# Nopp140-chaperoned 2’-O-methylation of small nuclear RNAs in Cajal bodies ensures splicing fidelity

**DOI:** 10.1101/2021.04.29.441821

**Authors:** Jonathan Bizarro, Svetlana Deryusheva, Ludivine Wacheul, Varun Gupta, Felix G.M. Ernst, Denis L.J. Lafontaine, Joseph G. Gall, U. Thomas Meier

## Abstract

Spliceosomal small nuclear RNAs (snRNAs) are modified by small Cajal body (CB) specific ribonucleoproteins (scaRNPs) to ensure snRNP biogenesis and pre-mRNA splicing. However, the function and subcellular site of snRNA modification are largely unknown. We show that CB localization of the protein Nopp140 is essential for concentration of scaRNPs in that nuclear condensate; and that phosphorylation by casein kinase 2 (CK2) at some 80 serines targets Nopp140 to CBs. Transiting through CBs, snRNAs are apparently modified by scaRNPs. Indeed, Nopp140 knockdown-mediated release of scaRNPs from CBs severely compromises 2’-O-methylation of spliceosomal snRNAs, identifying CBs as the site of scaRNP catalysis. Additionally, alternative splicing patterns change indicating that these modifications in U1, U2, U5, and U12 snRNAs safeguard splicing fidelity. Given the importance of CK2 in this pathway, compromised splicing could underlie the mode of action of small molecule CK2 inhibitors currently considered for therapy in cholangiocarcinoma, hematological malignancies, and COVID-19.

## INTRODUCTION

Spliceosomal small nuclear RNAs (snRNAs) are the work horses at the core of the nuclear splicing machinery multiplying and diversifying the protein coding potential of the genome (Wilkinson et al., 2019). In addition to ribosomal RNAs (rRNAs) and transfer RNAs (tRNAs), snRNAs constitute the most abundant non-coding RNAs. These abundant snRNAs of the major spliceosome are U1, U2, U4, U5, and U6. All but U6 transit through the cytoplasm for their assembly into functional small nuclear ribonucleoproteins (snRNPs) (Will and Lührmann, 2001; Yong et al., 2004). In the cytoplasm, they acquire a heptameric Sm protein ring and a tri-methyl cap while their 3’-end is trimmed. Upon re-entry into the nucleus, snRNAs are modified at specific ribose moieties by 2’-O-methyl groups and at specific bases by isomerization of uridines to pseudouridines (Bohnsack and Sloan, 2018; Karijolich and Yu, 2010). Additionally, they assemble into functional snRNPs with their remaining complement of snRNP-specific proteins. Although it is known that pseudouridines in snRNAs are essential for proper assembly into snRNPs and for pre-mRNA splicing (Yu et al., 1998; Zhao and Yu, 2004), the role of the 2’-O-methyl groups remains poorly defined. It has been particularly challenging to access the roles of snRNA modifications in intact cells. Here we uncover an explanation for their cellular functions.

The function of snRNA modifications in pre-mRNA splicing and spliceosome assembly has been of longstanding interest and has been assayed by multiple approaches (Bohnsack and Sloan, 2018). Thus, abundant and fully modified snRNAs could readily be purified and compared to in vitro transcribed, unmodified snRNAs. This was performed in snRNA depleted Xenopus oocytes and in cell-free splicing systems with yeast and HeLa cell nuclear extracts. In this manner, successful pre-mRNA splicing was observed with unmodified snRNAs U1, U4, U5, and U6 (Fabrizio et al., 1989; Ségault et al., 1995; Wersig and Bindereif, 1992; Will et al., 1996).

However, in vitro transcribed U2 snRNA was unable to support splicing pointing to the importance of the 14 pseudouridines and ten 2’-O-methylgroups in mammalian U2 snRNA (Pan and Prives, 1989; Ségault et al., 1995). Interestingly, analysis of in vitro transcribed U2 snRNA that supported splicing in yeast extracts revealed only pseudouridines but no 2’-O-methyl groups (McPheeters et al., 1989). Apparently pseudouridylation occurred during reconstitution providing evidence for the importance of pseudouridylation of U2 snRNA for the splicing reaction. We now document similar importance for two specific 2’-O-methyl groups in U2 snRNA.

The enzymes responsible for snRNA modifications have been identified, the small Cajal body (CB) specific RNPs (scaRNPs). Two major classes of scaRNPs are distinguished by their H/ACA and C/D scaRNAs, which guide pseudouridylation and 2’-O-methylation, respectively, of snRNAs by site-specific base pairing (Kiss, 2001; Maxwell and Fournier, 1995; Smith and Steitz, 1997). Each scaRNP consists of one of many distinct scaRNAs, of four core proteins, and of the CB-specifying protein WDR79 (aka TCAB1). In case of H/ACA scaRNPs, the four core proteins include the pseudouridine synthase NAP57 (aka dyskerin) and in case of C/D scaRNPs, the methyl transferase fibrillarin (Kiss, 2004; Meier, 2005). As the name suggests, scaRNPs are concentrated in CBs (Massenet et al., 2016; Meier, 2016). Similarly, snRNAs accumulate in and transit through CBs. Based on this confluence of the scaRNP enzymes and their target snRNAs in CBs, CBs have long been implicated as the sites of snRNA modification (Darzacq et al., 2002). Indeed, exogenous snRNA constructs are only modified when targeted to CBs but not to nucleoli (Jady 2003). We now certify CBs as the sites of endogenous snRNA modification.

CBs were discovered over 100 years ago and have been known by various names, accessory body of the nucleolus, coiled bodies, and now CBs, but their function is still shrouded in mystery (Gall, 2003; Machyna et al., 2013). Like many nuclear bodies, CBs are mainly defined by their composition. In addition to snRNPs and scaRNPs, CBs are highly enriched among other proteins in coilin, the CB-specific protein of still unknown function, in the survival of motor neuron protein SMN (which is required for the cytoplasmic assembly of heptameric Sm protein rings onto snRNAs), and the trimethyl guanosyl synthase TGS1 (which is responsible for the cytoplasmic hypermethylation of the snRNA monomethyl caps) (Raimer et al., 2017; Will and Lührmann, 2001). Confusingly, therefore, the nuclear CBs are enriched in enzymes with well-defined cytoplasmic functions. In addition to snRNA modification, CBs have been implicated in recycling of snRNPs during spliceosome assembly (Chen et al., 2017; Staněk and Neugebauer, 2006). Nopp140, the chaperone of scaRNPs and small nucleolar RNPs (snoRNPs), in addition to nucleoli, also concentrates in CBs (Meier, 2005; Meier and Blobel, 1994). In coilin knockout cells, residual CBs remain that harbor Nopp140 and scaRNPs but not snRNPs and SMN supporting a scaRNP-specific function for Nopp140 in CBs (Tucker et al., 2001). More recently, formation of CBs and nucleoli have been assigned to the mechanism of liquid-liquid phase separation (Brangwynne et al., 2011; Lafontaine et al., 2020; Zhu and Brangwynne, 2015).

Indeed, the enrichment in CBs of RNAs, RNA binding proteins, and proteins with intrinsically disordered domains, e.g., Nopp140 and coilin, provide fertile ground for such a mechanism (Meier and Blobel, 1994; Na et al., 2018; Neugebauer, 2017; Tantos et al., 2013).

Small CB-specific RNAs were originally identified as snoRNAs in nucleoli. In fact, consisting of box H/ACA and C/D RNAs and the four box-specific core proteins, snoRNPs are the most abundant guide RNPs. They pseudouridylate and 2’-O-methylate some 100 nucleotides each of mammalian pre-rRNA by site-directed hybridization in nucleoli. Each sno/scaRNA has the potential to guide modification of two nucleotides in their target RNAs. Although sno- and scaRNPs overall exhibit some clear differences in substrates, subcellular location, and even the additional WDR79 protein of scaRNPs, the lines are blurred because some scaRNAs exhibit complementarity to both snRNA and rRNA (Kiss et al., 2004; Meier, 2016). Therefore, some sno/scaRNPs have the potential to function in both nucleoli and CBs. Interestingly, the only known none-RNP protein found in both organelles is Nopp140.

Nucleoli, the sites of ribosome synthesis and assembly, are by far the most phase-dense bodies of every cell. Their formation is apparently aided by liquid-liquid phase separation (Brangwynne et al., 2011; Lafontaine et al., 2020; Zhu and Brangwynne, 2015). By transmission electron microscopy, three distinct regions are discernable, fibrillar centers (FCs) and the surrounding dense fibrillar component (DFC), which are altogether embedded in the granular component (GC). The rRNA genes are located in the FCs and transcribed at the border to the DFC into which the nascent rRNAs extend while being modified and processed (Derenzini et al., 1990; Dundr and Misteli, 2001; Hadjiolov, 1985; Scheer and Hock, 1999; Spector, 1993). Many subsequent steps of maturation and assembly with ribosomal proteins occur in the GC, where various forms of pre-ribosomes are found before export to the cytoplasm. The home of snoRNPs are the most densely packed parts of nucleoli, the DFCs. Multiple factors are required in DFCs to orchestrate the dance between snoRNPs and nascent rRNAs while avoiding RNA tangles and electrostatic repulsion. Nopp140 and helicases are only some of such chaperones.

Nopp140 is a nucleolar (DFC) and Cajal body phosphoprotein encoded by the gene NOLC1 (Meier and Blobel, 1990, 1994). Except for its association with sno- and scaRNPs, its function is poorly defined. Although its N- and C-termini are evolutionarily most highly conserved, e.g., the last 50 amino acids of human Nopp140 exhibit 59% sequence identity with its yeast ortholog Srp40p, the ∼500 amino acid long central repeat domain is its most outstanding hallmark (Meier, 1996; Meier and Blobel, 1992). The repeat domain contains ten acidic serine stretches alternating with lysine, alanine, and proline-rich tracts. Most, if not all of the 82 serines of the acidic repeats are phosphorylated by casein kinase 2 (CK2) effecting a shift in migration on an SDS polyacrylamide gel of 40kD (Li et al., 1997; Meier and Blobel, 1992). This phosphorylation is required for the association of Nopp140 with sno- and scaRNPs, but not for their modification function (Wang et al., 2002). We now show that the phosphorylation of Nopp140 is required for its accumulation in CBs.

High-resolution CRISPR screens identified Nopp140 as essential for cell survival (Hart et al., 2015; Wang et al., 2015). Using a targeted CRISPR/Cas9 approach in polyploid HeLa cells, we established three cell lines with very low levels of Nopp140 (∼1-7% residual protein level), i.e., Nopp140 knockdown (KD) cell lines (Bizarro et al., 2019). Surprisingly, Nopp140 KD cells do not exhibit any growth or gross phenotypes. Nevertheless, the KD cells reveal subtle but clear differences in Nopp140 chaperoned activities filtering nonessential from essential functions. We showed that one of these nonessential functions is corralling scaRNPs in Cajal bodies (Bizarro et al., 2019). In Nopp140 low-expressing cells, all scaRNPs are released from Cajal bodies but the overall levels and integrity of the RNPs remain unaffected. As one of the consequences, the telomerase scaRNP is no longer sheltered in CBs but has continuous access to telomeres extending them gradually (Bizarro et al., 2019). Here we present the consequences of Nopp140 KD for all other scaRNPs when no longer maintained in CBs and for snoRNPs in nucleoli.

## RESULTS

### Establishment of stable Nopp140 rescue cells

In a prior study, we generated three stable Nopp140 knockdown (KD) cell lines, KD1a, KD1b, and KD2 originating from two HeLa parent lines P1 and P2 (Bizarro et al., 2019). In the Nopp140 KD cells, intact scaRNPs were displaced from CBs. This phenotype could be rescued by transient re-expression of Nopp140 establishing that it was not an off-target effect of our CRISPR/Cas9 approach (Bizarro et al., 2019). To allow for biochemical and genome-wide approaches of Nopp140 rescue, we reintroduced Nopp140 on a plasmid with a selectable marker into the Nopp140 KD2 cells followed by antibiotic resistance selection of single clones to obtain three stable rescue cell lines, Nopp140 R2a, R2b, and R2c. Indirect immunofluorescence localized Nopp140 and NAP57, the pseudouridine synthase of H/ACA RNPs, in nucleoli and CBs (arrows) in the P2 parent cells (Fig. 1A top). In contrast, in the Nopp140 KD2 knockdown cells, Nopp140 was lost from CBs and nucleoli whereas NAP57 was present in nucleoli but lost from CBs (Fig. 1A middle). Nopp140 R2a rescue cells uniformly expressed Nopp140 in both nucleoli and CBs (arrows) and rescued the CB localization of NAP57 (Fig. 1A bottom).

**Figure 1.**
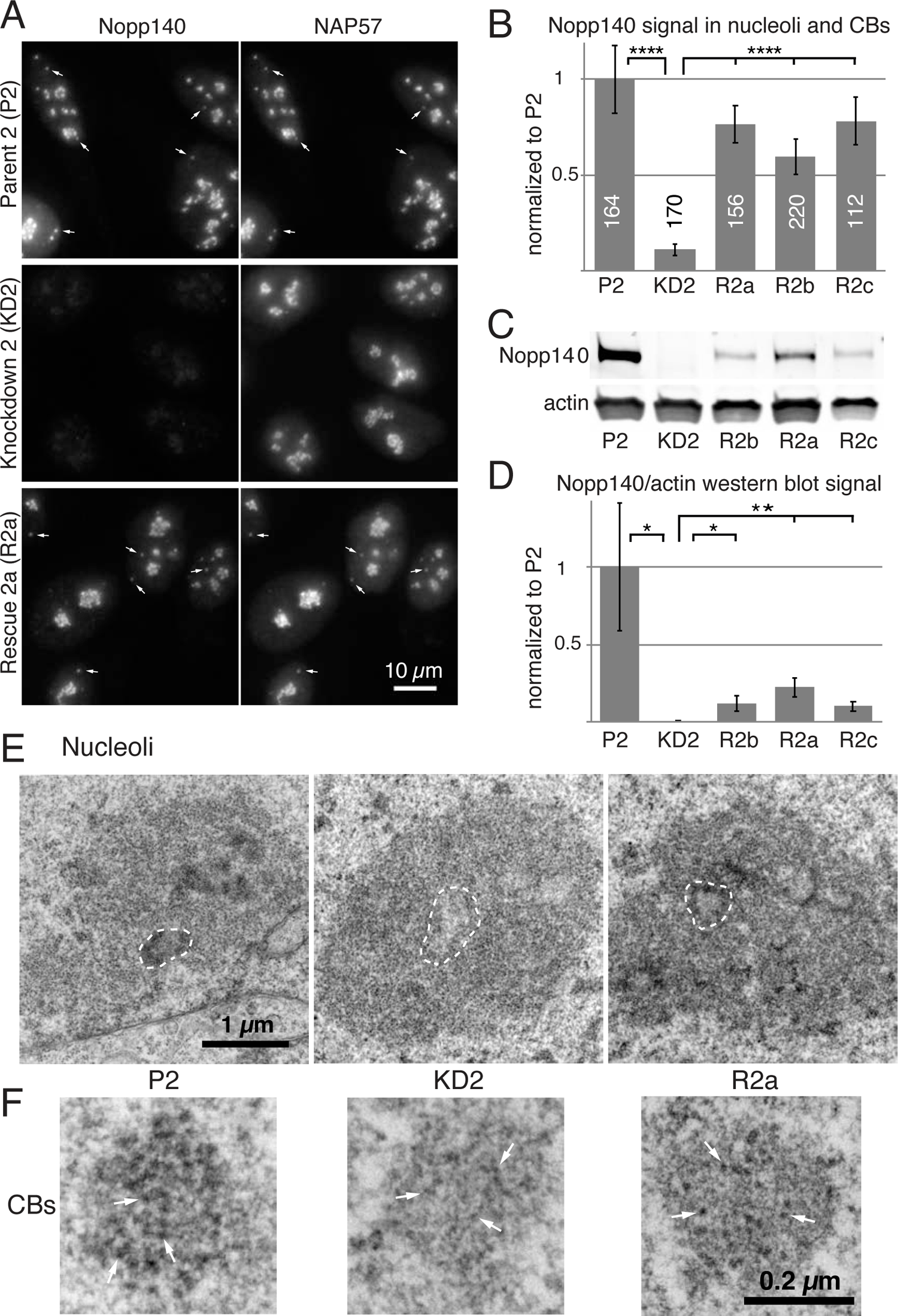
Effects of Nopp140 knockdown (KD) on nucleoli and Cajal Bodies (CBs) are restored in cells stably re-expressing Nopp140. (A) Indirect immunofluorescence with Nopp140 (left) and NAP57 antibodies (right) on parent 2 (P2; top), KD2 (middle), and rescue 2a cells (R2a; bottom). Note, 5 CBs are highlighted in the top and bottom panels (arrows). Neither Nopp140 nor NAP57 are visible in CBs (middle panels) and Nopp140 is strongly reduced in nucleoli (left). Scale bar = 10µm. (B) Quantification of Nopp140 fluorescent signal in nucleoli and CBs of P2, KD2, and rescue cells R2a-c normalized to P2 cells. The means ± standard deviations (SDs) are indicated. The numbers refer to the number of cells analyzed for each cell line. Unpaired t-tests identify significant differences (**** P < 0.0001). (C) Western blots of whole cell extracts of the same set of cells probed for Nopp140 (top) and actin (bottom) and detected by a near-infrared imaging system (Odyssey). (D) Quantification of the near-infrared signals of triplicate western blots shown in (C) and normalized to P2 signals. Means ± SDs are shown. Unpaired t-tests identify significant differences (* P < 0.05, ** P < 0.005). (E) Transmission electron micrographs of P2 (left), KD2 (center), and R2a (right) cells. One fibrillar center (FC) surrounded by a dense fibrillar component (DFC) is outlined in each nucleolus (dashed line). Note the striking loss of contrast of the DFC in the KD2 and its reappearance in the R2a cells re-expressing Nopp140. Scale bar = 1µm. (F) Transmission electron micrographs of one CB from each of the same cells as in (E). Three electron-dense granules are pointed out in each CB (arrows). Note the disappearance/shrinking of the granules in KD2 cells and their reappearance in R2a cells. Scale bar = 0.2µm.

According to fluorescent signal in nucleoli and CBs, all three rescue cell lines re-expressed Nopp140 to 60-80% of the parent cells (Fig. 1B). Surprisingly, when protein levels of Nopp140 in the rescue cells were assessed by western blotting, Nopp140 re-expression appeared more subtle (Fig. 1C). Apparently, the different dynamic range of the two immunodetection methods is responsible for this discrepancy. This is supported by the fact that Nopp140 re-expression was increased over 13-fold when assessed by western blotting (Fig. 1D, compare R2a-c to KD2) but only 7-fold when detected by indirect immunofluorescence (Fig. 1B).

Light microscopy did not detect any morphological differences between Nopp140 parent and KD cells, but alterations were noticed at the ultrastructural level (Bizarro et al., 2019). Differential contrast by electron microscopy identifies the classic tripartite structure of nucleoli, the light fibrillar centers (FCs) surrounded by the dense fibrillar component (DFC) that are altogether embedded in the granular component (GC) (Fig. 1E, P2, one FC-DFC unit is outlined). In the Nopp140 KD2 cells, the contrast of the DFC, where Nopp140 and snoRNPs reside, was lost (Fig. 1E, KD2), but reappeared in the Nopp140 rescue cells (Fig. 1E, R2a). In case of CBs, the loss of scaRNPs caused a marked reduction in contrast and a halving in size of the granules making up their coils (Fig. 1F, compare KD2 to P2). In contrast, CB granule size and contrast in the rescue cells was mostly restored (Fig. 1F, R2a). Together with the repopulation of CBs with scaRNPs in the rescue cells, this data further indicates that scaRNPs normally reside in the granules of CBs. We previously reported the same effects of Nopp140 KD on nucleoli and CBs in other KD cells, KD1a, demonstrating that this is not a clonal aberration (Bizarro et al., 2019). Throughout this study, we now employ three Nopp140 rescue cell lines to document that the observed effects are not due to some unrelated event but specifically to Nopp140 depletion.

**Figure 2.**
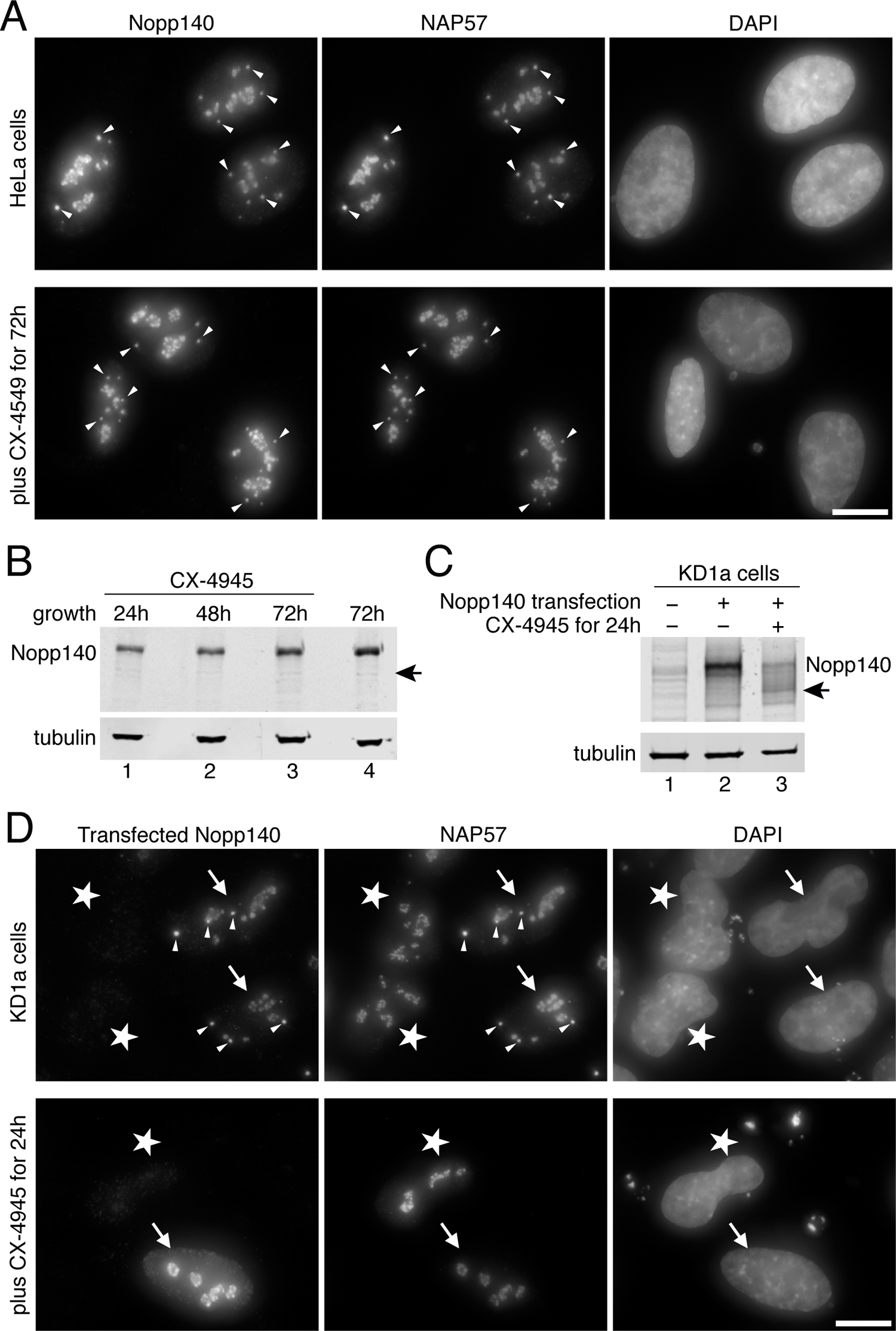
Nopp140 phosphorylation by casein kinase 2 (CK2) is required for CB localization. (A) Top panels: indirect immunofluorescence for Nopp140 (left) and NAP57 (center) with DAPI DNA stain (right) on control P2 cells (top). Bottom panels: the same, but after a 72h incubation with 10µM concentration of the CK2 inhibitor CX-4945 (silmitasertib). Note, both proteins remain in nucleoli and CBs under both conditions. Some CBs are highlighted (arrowheads). Scale bar = 10µm. (B) Western blots on whole P2 cell lysates as described for Fig. 1C after incubation for various number of hours (top) with 10µM CX-4945 (lanes 1-3) and without (lane 4). The migrating positions of Nopp140 and tubulin are indicated (left), and that of unphosphorylated Nopp140 (right, arrow). (C) Western blots of KD1a cell extracts after 24h transient transfection with Nopp140 (lanes 2 and 3) or untransfected (lane 1), with (lane 3) and without (lanes 1 and 2) 10µM CX-4945. Labeling as in (B). (D) Top panels: indirect immunofluorescence of KD1a cells transfected (arrows) or untransfected (asterisks) with Nopp140 and stained for Nopp140 (left) and NAP57 (center) with DNA stain (right). Note transfected Nopp140 localizes to both nucleoli and CBs in these Nopp140 knockdown cells, whereas endogenous NAP57 localizes to nucleoli independent of the presence of Nopp140 but to CBs only in Nopp140 transfected cells (some CBs are highlighted by arrowheads). Bottom panels: same as top but in the presence of the CK2 inhibitor CX-4945. Note, newly synthesized and unphosphorylated Nopp140 accumulates only in nucleoli but not CBs and accordingly NAP57 is not recruited to CBs either. Scale bar = 10µM.

### Phosphorylation of Nopp140 is required for CB localization

The most outstanding feature of the Nopp140 amino acid sequence is its 10 alternating acidic serine and positively charged lysine-, alanine-, and proline-rich repeats (Meier and Blobel, 1992). Casein kinase 2 (CK2) is responsible for the phosphorylation of some 80 serines in the 10 acidic serine stretches effecting a 40kD shift in migration on denaturing polyacrylamide gels (SDS-PAGE) (Meier, 1996; Meier and Blobel, 1992). It is this extreme phosphorylation of Nopp140 that forms the basis for its interaction with sca- and snoRNPs because dephosphorylated Nopp140 no longer associates with the RNPs (Wang et al., 2002). Given that scaRNPs are lost from CBs concomitantly with Nopp140, we tested if phosphorylation of Nopp140 by CK2 is required for CB accumulation. We first tested if simple inhibition of CK2 affected localization of Nopp140. We used the small molecule CX-4945 (silmitasertib), a selective ATP-competitive inhibitor of CK2 (Siddiqui-Jain et al., 2010). Even after three days of incubation with the inhibitor, when most cells had died, Nopp140 and NAP57 localization was unaffected in the surviving cells where they remained in CBs and nucleoli (Fig. 2A, lower panels). To ascertain that Nopp140 lost its phosphorylation during the incubation period, migration of Nopp140 on SDS-PAGE was analyzed by western blotting (Fig. 2B). Surprisingly, at all time points of CK2 inhibition, Nopp140 migrated at 140kD and not at the 100kD characteristic for the unphosphorylated protein (Fig. 2B, arrow). This indicated that the phosphorylation of Nopp140 did not turn over while the CK2 inhibitor caused cell cycle arrest (Siddiqui-Jain et al., 2010). Hence, we tested if the CK2 inhibitor prevented only phosphorylation of newly synthesized Nopp140. Taking advantage of our Nopp140 KD cells, which express little to no Nopp140 (Fig. 2C, lane 1), Nopp140 was transiently transfected in the absence and presence of the CK2 inhibitor CX-4945. The phosphorylation status of Nopp140 was assessed by its migration on SDS-PAGE. Thus, newly synthesized and fully phosphorylated Nopp140 was only detected in the absence of CK2 inhibitor (Fig. 2C, lane 2). However, the inhibitor mostly prevented Nopp140 phosphorylation, which migrated at its unphosphorylated position of 100kD (Fig. 2C, lane 3). Additionally, some more slowly migrating bands, representing intermediate degrees of phosphorylation, were observed. Therefore, while leaving already phosphorylated Nopp140 untouched, the CK2 inhibitor effectively prevented phosphorylation of newly synthesized Nopp140. To investigate the localization of Nopp140 and NAP57 under these conditions, we performed indirect immunofluorescence experiments (Fig. 2D). In the absence of the CK2 inhibitor, transfection of Nopp140 (cells with arrows) caused both proteins to localize to CBs and nucleoli. However, in the residual untransfected Nopp140 KD cells (asterisks), NAP57 localized only to nucleoli but not CBs (Fig. 2D, upper panels) consistent with our previous results (Bizarro et al., 2019). In the presence of CX-4945, newly translated, unphosphorylated Nopp140 similarly accumulated only in nucleoli but not CBs (Fig. 2D, lower panels, arrow) demonstrating that Nopp140 phosphorylation was required for CB targeting. Consistent with the fact that NAP57 accumulation in CBs depends on the localization of Nopp140 in CBs (Bizarro et al., 2019), NAP57 stayed in nucleoli but was excluded from CBs in the presence of CX-4945, even in Nopp140 transfected cells (Fig. 2D, lower panels, arrow). In summary, CK2 phosphorylation of Nopp140 is required for the accumulation of both proteins in CBs and by extension for that of scaRNPs.

Thus, the molecular consequences of displacement of scaRNPs from CBs, described in the remainder of this manuscript, need to be considered in CX-4945 (silmitasertib) therapy. Silmitasertib is currently in a phase II trial of cholangiocarcinoma (NCT02128282) as well as being considered as a drug against hematological malignancies and COVID-19 (Bouhaddou et al., 2020; Chon et al., 2015; Silva-Pavez and Tapia, 2020). As outlined below, compromised splicing fidelity due to reduced 2’-O-methylation of snRNAs may thus contribute to the molecular mode of action of this CK2 inhibitor.

### ScaRNP depletion from CBs alters snRNA modification

Due to the conspicuous colocalization in CBs of scaRNP enzymes and their snRNA targets, CBs have long been presumed the subcellular sites of snRNA modification, but this was never demonstrated for endogenous particles in mammalian cells (Darzacq et al., 2002; Deryusheva et al., 2012; Jády et al., 2003). After having documented the consequences of telomerase displacement from CBs, we investigated the consequence of displacement of all other scaRNPs from CBs. In Nopp140 KD cells, scaRNPs remained intact and their cellular levels unaltered indicating that the bulk of scaRNPs in CBs separated from their snRNP targets and dispersed in the nucleoplasm (Bizarro et al., 2019). We thus asked if and to what extent snRNAs were still modified in Nopp140 KD cells.

We started with analysis of U2, which with 14 pseudouridines and ten 2’-O-methyl groups is the most highly modified spliceosomal snRNA (Morais et al., 2021). In fact, one quarter of its 5’-half nucleotides are modified. The modifications are important for snRNP biogenesis and pre-mRNA splicing (Dönmez et al., 2004; Yu et al., 1998; Zhao and Yu, 2004). To map modified residues of snRNAs, we isolated total RNA from parent and Nopp140 KD cells and performed several reverse transcriptase-based assays after chemical treatment of the RNA with CMC [N-cyclohexyl-N’-(2-mopholinoethyl) carbodiimide metho-p-toluene sulfonate] to identify pseudouridines and in the presence of low dNTP concentrations to recognize 2’-O-methyl groups (Bakin and Ofengand, 1993; Maden et al., 1995). Under these conditions, strong stops are observed during RT revealing pseudouridines and 2’-O-methylated residues (Fig. 3A-E).

**Figure 3.**
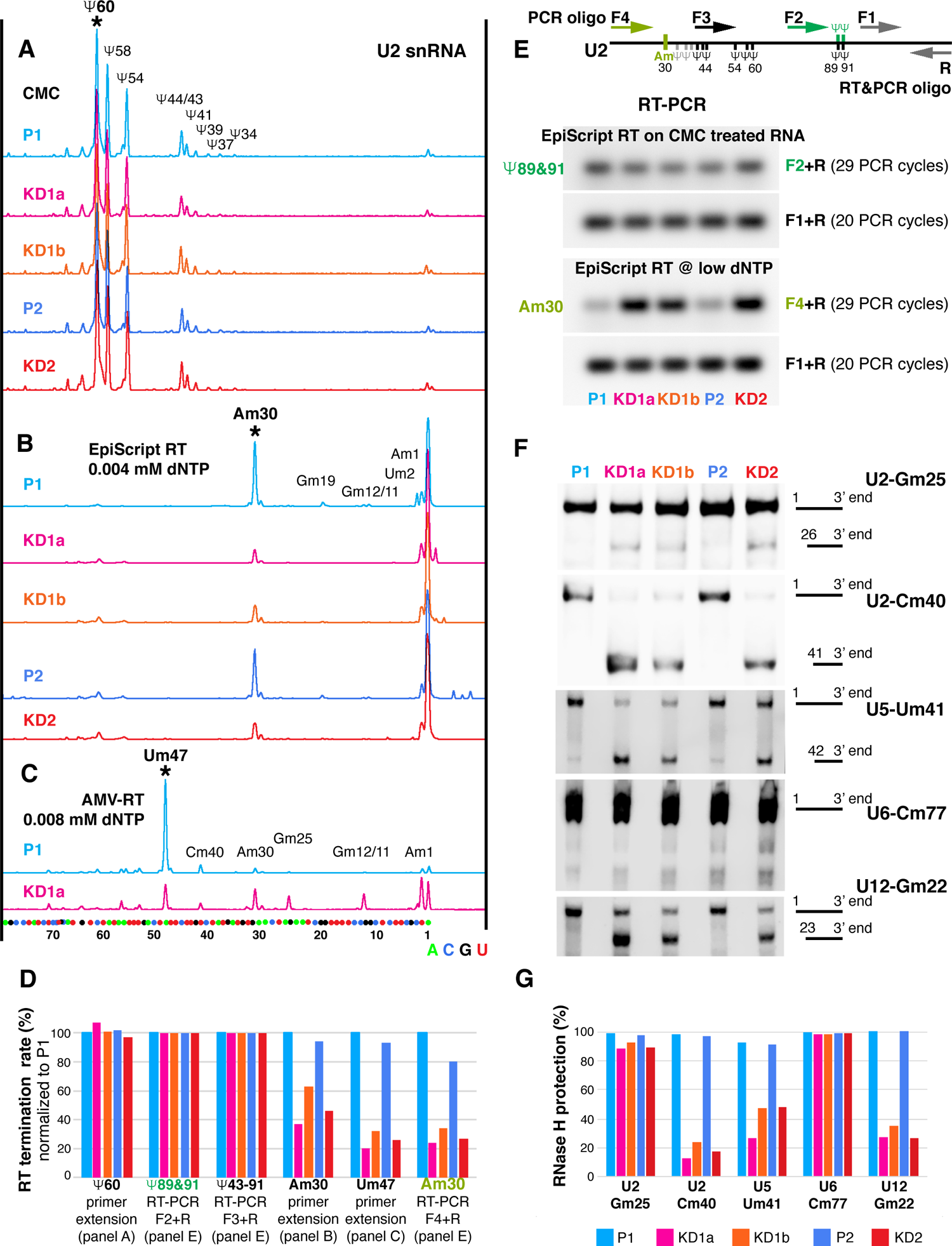
2’-O-methylation, but not pseudouridylation, is reduced at most sites of RNA polymerase II transcribed snRNAs. (A-C) Mapping of modified nucleotides in U2 snRNA of Nopp140 parent (P1 and P2; blue tones) and knockdown cells (KD1a, KD1b, and KD2; red tones) using fluorescent primer extension. Nucleotide positions were aligned to sequencing reactions on in vitro-transcribed U2 snRNA (bottom). (A) Nine pseudouridines are detectable through strong stops after CMC treatment. Note the same height of all peaks in total RNAs isolated from all five cell lines, indicating undisturbed pseudouridylation of U2 snRNA after Nopp140 KD. (B) The 2’-O-methylated residue Am30 of U2 is readily detectable through a strong stop using EpiScript RT at low dNTP concentration in the parent traces (blue tones) but is severely reduced in the KD traces (red tones). Due to the initial strong stop at Am30, the subsequent 2’-O-methylated residues could not be determined reliably. (C) The 2’-O-methylated residue Um47 is identified by a strong stop using AMV-RT at low dNTP levels in U2 from P1 (blue) and is severely reduced in KD1 cells (pink). Subsequent 2’-O-methylated residues are masked due to the strong initial stop. (D) Quantification of RT termination rates normalized to P1, i.e., the degree of modification, of the indicated residues in panels A to C (peaks marked by asterisks) and in panel E (green and olive). Additionally, the termination rates across 7 Ψs of U2 snRNA (Ψ43-91) are quantified (primers F3 and R1 in schematic of panel E). Note the greater reduction in KD1a (pink) over KD1b (orange) mirroring the residual Nopp140 levels and the lack of effect on any of the Ψs tested (key in panel G). (E) Semi-quantitative RT-PCR of U2 using EpiScript RT on CMC derivatized RNA with primers F2 and R1 to detect Ψ89 and Ψ91 (green) and at low dNTP concentration with primers F4 and R1 to detect 2’-O-methylated Am30 (olive). The amplification scheme is depicted on top and the gels of the PCR products are below. Note, EpiScript RT does not detect 2’-O-methylated residues upstream of Am30 (B). RNA from all cell lines was analyzed and the results expressed relative to the amplification products of an unmodified stretch of U2 (primers F1 and R) are quantified in (D). (F) RNase H protection assays to quantitatively detect the degree of 2’-O-methylation of snRNAs from all parent and KD cell lines at U2-Gm25, U2-Cm40, U5-Um41, U6-Cm77, and U12-Gm22. Note, the more full-length products are detected the more the specific site is 2’-O-methylated. (G) Percent of RNase H protection for each cell line and nucleotide in (F).

Using fluorescent primer extension on CMC-derivatized U2 snRNA, we first mapped pseudouridines (Fig. 3A). As the peak size of none of the 9 pseudouridines varied significantly between the two parent cells (Fig. 3A, blue) and the three knockdown cells (Fig. 3A, red), we conclude that pseudouridylation of U2 snRNA was unaffected by Nopp140 KD. Quantification of the first strong stop marking the pseudouridine at residue 60 of U2 snRNA (Fig. 3A, Ψ60) confirmed the same degree of pseudouridylation of parent and KD cells (Fig. 3D). Using an independent assay, we globally mapped the pseudouridines of CMC-derivatized U2 snRNA by semi-quantitative RT-PCR (Fig. 3E, upper panel, green). Indeed, RT-PCR across Ψ89 and Ψ91 and a stretch including 7 Ψs of U2 snRNA (Ψ43-91) confirmed the same RT termination rate between the parent and Nopp140 KD cells relative to that of a 3’ unmodified stretch (Fig. 3E, F3/F2 relative to F1). Quantification of the amplified bands normalized to U2 snRNA from the parent cells confirmed our conclusions (Fig. 3D). In summary, U2 snRNA is fully pseudouridylated, even in the absence of scaRNP pseudouridylases from CBs where U2 snRNA remains concentrated (Bizarro et al., 2019). Apparently, pseudouridylation of snRNAs is too important to be lost and instead occurs in the nucleoplasm of Nopp140 KD cells.

Next, we assessed the degree of ribose methylation of U2 snRNA. This was performed using low dNTP concentrations during reverse transcription with two different reverse transcription enzymes (RTs) that show differential sensitivity towards individual sites of 2’-O-methylation. In contrast to pseudouridines, the levels of all mapped 2’-O-methyl groups were significantly reduced in U2 snRNA isolated from KD cells (Fig. 3B and D, blue traces) relative to those from parent cells (traces in red tones). Quantification of the termination rate at Am30 (Fig. 3B) and Um47 (Fig. 3C) verifies the reduction of 2’-O-methylation at those sites of U2 snRNA in Nopp140 KD versus parent cells (Fig. 3D). We further confirmed the results for U2-Am30 using an independent semi-quantitative RT-PCR assay using EpiScript RT at low dNTP concentration, which is mainly sensitive to the 2’-O-methyl group at A30 but not those further downstream (Fig. 3E, lower panels). The amplification efficiency across Am30 (Fig. 3E, lower panels, F4+R) was compared to that of an unmodified stretch of U2 further downstream (F1+R). Normalized quantification of the results for all parent and KD cells corroborated those obtained through fluorescent primer extension analysis (Fig. 3D). Remarkably, the levels of 2’-O-methylation at Am30 and Um47 mirrored the degree of Nopp140 KD in those cells (Bizarro et al., 2019), i.e., lowest levels were detected in U2 snRNA from KD1a and KD2 cells, whereas intermediate levels were detected in those from the KD1b cells (Fig. 3D). Thus, the loss of 2’-O-methylation at U2-Am30 and Um47 correlated with the loss of Nopp140 and scaRNPs from CBs indicating that this snRNA modification normally occurs in CBs and is not fully required under standard conditions for growth in our cell lines.

To test if the loss of 2’-O-methylation was specific to U2 snRNA, we used a quantitative site-specific assay on U5-Um41, U6-Cm77, and U12-Gm22, in addition to U2-Gm25 and U2-Cm40. Specifically, we employed RNase H mediated site-specific cleavage with chimeric RNA/DNA oligos (Fig. 3F). In this assay, cleavage of RNA is prevented when the ribose at the specific residues is 2’-O-methylated (Yu et al., 1997). In the Nopp140 KD cells KD1a, KD1b, and KD2, methylation of U2-Cm40, U5-Um41, and U12-Gm22 was mostly lost, i.e., cleavable (Fig. 3F). In contrast, in the parental cell lines P1 and P2, no observable cleavage was noted at those nucleotides indicating complete 2’-O-methylation (Fig. 3F). Remarkably, 2’-O-methylation at one residue of U2 (Gm25) and of U6 (Cm77) was unaffected in all cells (Fig. 3F). In the case of U6 snRNA this was not surprising because its 2’-O-methylation guide RNPs concentrate in nucleoli and not CBs (Ganot et al., 1999). The level of RNase H protection at these 5 sites of the 4 snRNAs was quantified (Fig. 3G). Importantly, the loss of 2’-O-methylation was due to that of Nopp140 because the modification of U2-Cm40, U2-Cm61, U5-Um41, and U12-Gm22 was fully restored in Nopp140 re-expressing cell lines (Supplemental Table S1). Additionally, the degree of loss of methylation again mirrored that of Nopp140 (Bizarro et al., 2019), i.e., more loss from KD1a than KD1b snRNAs (Fig. 3G). Finally, methylation at G25 of U2 snRNA seems too vital to be lost and apparently occurs in the nucleoplasm of Nopp140 KD cells.

### RiboMethSeq captures most 2’-O-methylation sites in snRNAs

To corroborate and expand our findings of changes in 2’-O-methylation of stable RNAs in Nopp140 KD cells, we used the systematic mapping approach of RiboMethSeq (RMS). This approach takes advantage of differential sensitivity to alkaline cleavage of 2’-O-methylated versus unmethylated residues that becomes statistically apparent during deep sequencing of alkaline cleaved abundant RNAs (Birkedal et al., 2015; Krogh et al., 2016; Marchand et al., 2016; Sharma et al., 2017). This method is particularly efficient in identifying the degree of 2’-O-methylation in rRNA but, as described below, also allows monitoring these modifications in other abundant RNAs, such as snRNAs (Krogh et al., 2017).

In fact, with a sequencing depth of 25 million reads per sample, RMS was sufficiently sensitive to reliably identify most 2’-O-methylated residues in the major spliceosomal snRNAs U1, U2, U5, and U6 (Fig. 4 and Supplemental Table S1). However, coverage of U4 was insufficient to provide statistically significant scores in our sequencing. The RMS score (fraction of a residue that is 2’-O-methylated) of most residues of snRNAs from the parental P2 cell lines was equal to or above 0.7 (Fig. 4A, blue, and Supplemental Table S1). In contrast, the RMS scores for most residues of U1, U2, and U5 snRNAs from the Nopp140 KD2 cells were below 0.7 (Fig. 4A, red, and Supplemental Table S1) yielding ratios of KD2 over P2 of below 0.6 indicating a significant reduction in modification of those residues (Fig. 4B). In contrast, the KD2/P2 ratios of all five 2’-O-methylated residues of U6 snRNA were above 0.7, indicating that 2’-O-methylation of U6 was not or barely affected by Nopp140 KD. This confirmed our RNase H cleavage-based results for U6-Cm77 (Fig. 3F and G) and supports U6 modification occurring outside CBs (Deryusheva and Gall, 2019; Ganot et al., 1999). Interestingly, in contrast to 5 other U2 residues, 2’-O-methylation of U2-G12 and U2-G25 was barely impacted by Nopp140 KD (Fig. 4A-C). For U2-Gm25, this lack of effect on 2’-O-methylation was also noted by RNase H mediated cleavage with RNA/DNA hybrid oligonucleotides in all three Nopp140 KD cell lines (Fig. 3F and G, and Supplemental Table S1). The robustness of these results was confirmed by an independent second round of RMS (Fig. 4C, P2‘/KD2’). Importantly, the loss of methylation at all snRNA sites was restored in our rescue cells establishing it as a consequence of Nopp140 KD (Fig. 4C, R2a’). All individually determined and RMS scores of snRNA modification agree with each other and are numerically summarized in Supplemental Table S1. Apparently, the 2’-O-methylation of U2-Gm12 has to be added to that of U2-Gm25 as too vital to be lost or being mediated by a yet to be identified chaperone (Fig. 4A-C and Supplemental Table S1). We conclude that the loss of scaRNPs from CBs impacts 2’-O-methylation, but not pseudouridylation, of snRNAs transiting through these condensates.

**Figure 4.**
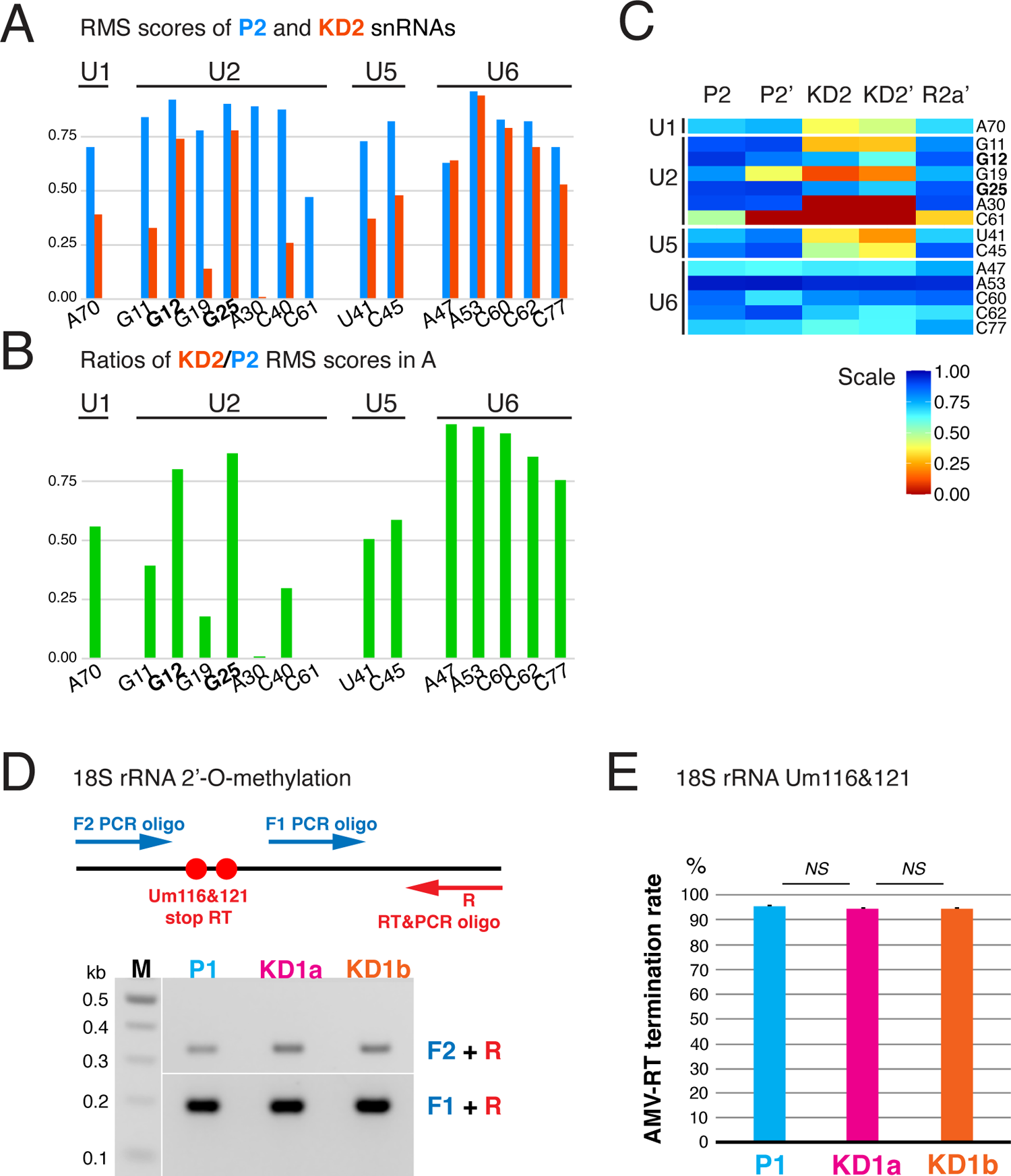
RiboMethSeq (RMS) shows a severe reduction of 2’-O-methylation at most sites of U1, U2, and U5 snRNAs but not U6 (A-C), nor at Um116 and 121 of 18S rRNA detected by RT-PCR (D-E). (A) Histogram of RMS scores for U1, U2, U5, and U6 snRNAs obtained from RMS of total RNA isolated from Nopp140 parent (P2, blue) and knockdown (KD2, red) cells. Note the reduction in 2’-O-methylation at all sites except those in U6 and at Gm12 and Gm25 of U2 snRNA (bold). (B) Histogram of the KD2/P2 ratios of the RMS scores in (A). (C) Heat map of RMS scores of snRNAs from two experiments on total RNA isolated on separate occasions from P2, KD2, and rescue cells (R2a). The apostrophe indicates a separate experiment. Note the remarkable agreement of the RMS scores from the two experiments (P2 vs. P2’ and KD2 vs. KD2’) and the rescue of the loss of 2’-O-methylation after re-expression of Nopp140 (R2a’). (D) Semi-quantitative RT-PCR of 18S rRNA at low dNTP concentration with primers F2 and R relative to unmodified rRNA with primers F1 and R to detect 2’-O-methylated Um116&121 on P1 and KD1a and b cell RNA. Schematic on top and gel of products underneath. (E) Histogram of percent of AMV-RT termination rates of triplicated experiments expressed as the ratio of intensities of the two bands in D corrected for fragment length (mean ± SD).

### 2’-O-Methylation of rRNA remains mostly unaffected

In addition to scaRNPs in CBs, Nopp140 also associates with snoRNPs in nucleoli. There, snoRNPs are responsible for the modification of rRNA. In the absence of any obvious impact on ribosome synthesis, one of the most remarkable hallmarks of the Nopp140 KD cells is the reorganization of nucleoli evidenced by loosening or loss of contrast of the nucleolar DFC (Fig. 1E, KD2) (Bizarro et al., 2019). It is in this compartment where Nopp140 and all snoRNPs normally concentrate and modify nascent pre-ribosomal RNA guided by site-directed base pairing. To assess the impact on rRNA 2’-O-methylation, we first employed our semi-quantitative RT-PCR approach described above for U2 snRNA (Fig. 3E). Specifically, methylation at residues U116 and U121 of 18S rRNA was investigated in our Nopp140 parental P1 and KD1a and b cell lines (Fig. 4D). No difference in amplification efficiency was noted between all three RNA sample templates and across the methylated region versus an unmodified stretch of 18S rRNA (Fig. 4D, bottom). Quantification of the AMV-RT termination rates confirmed that Nopp140 KD had no impact on 18S rRNA methylation at residues Um116 and Um121 (Fig. 4E). This result indicated that at the tested positions 2’-O-methylation of rRNA, unlike that of snRNA, remained unaffected by Nopp140 KD.

To systematically survey the impact of Nopp140 KD on pre-rRNA 2’-O-methylation at each position known to be modified, we next employed RMS. Previous studies reported 2’-O-methyl modification scores of rRNA for different cancer cell lines under various conditions including transient KD of fibrillarin, the methyltransferase of C/D snoRNPs (Erales et al., 2017; Incarnato et al., 2016; Krogh et al., 2016; Sharma et al., 2017; Taoka et al., 2018). To assess the impact of Nopp140 KD on rRNA methylation, we considered in duplicates the ratio of RMS scores of the knockdown (KD2 and KD2’) over the parental (P2 and P2’) cell rRNAs. The ratios of most modified residues were above 0.8 (Supplemental Table S2, framed red). To ascertain a true effect, we used an extra margin by considering a ratio of ≤ 0.7 as reduced. Thus, out of 38 robustly modified residues of 18S rRNA in all studies, only 5 residues (13%) were affected by Nopp140 KD (Supplemental Table S2, 18S colored, framed red). All but one (Supplemental Table S2, 18S orange) of those 5 residues were constitutively hypomodified (RMS score ≤ 0.7) in at least one of the prior studies (Supplemental Table S2, 18S yellow) (Erales et al., 2017; Krogh et al., 2016; Sharma et al., 2017; Taoka et al., 2018). Out of the 65 robustly modified residues of 28S rRNA in all studies, 2’-O-methylation of 12 residues (18%) was reduced in Nopp140 KD cells (Supplemental Table S2, 28S colored, framed red). All but two of those 12 were constitutively hypomodified or, in the case of Cm2422, the RMS score ratio was significantly reduced (≤ 0.7) after fibrillarin KD in at least one of the prior studies (Supplemental Table S2, 28S yellow) (Erales et al., 2017; Krogh et al., 2016; Sharma et al., 2017; Taoka et al., 2018). Only 3 of the residues identified as hypomodified in most studies were unaffected by Nopp140 KD (Supplemental Table S2, 28S, green, Gm1316, Cm1881, and Um2415).

Nevertheless, most of the rRNA modifications that were reduced after Nopp140 KD corresponded to hypomodified or naturally sensitive sites of methylation (Erales et al., 2017; Krogh et al., 2016; Sharma et al., 2017). Given the normal proliferation of those cells, this suggested that these sites may be less important for ribosome assembly and function. However, the most interesting changes in rRNA methylation are those three residues in 18S and 28S rRNA that are normally fully modified in all studies, even under transient fibrillarin depletion, i.e., 18S-Um428, 28S-Gm4370, and 28S-Cm4456 (Supplemental Table S2, orange). Apparently, the chaperoning role of Nopp140 is particularly critical for those snoRNPs that are responsible for guiding methylation at those sites. A heatmap representation of the RMS score tables for all rRNAs visually confirms the above points (Fig. 5A). The affected residues are marked with yellow and orange dots and the unaffected but hypomodified residues are indicated with green dots following the color scheme of Supplemental Table S2 (Fig. 5A). The heatmap further underscores the reproducibility of the two parent (P2 and P2’) and knockdown RMS scores (KD2 and KD2’). Importantly, Nopp140 re-expression restored the levels of 2’-O-methylation at the affected sites of rRNA to those of parent cells (Fig. 5A, R2a’). Unsupervised clustering of the RMS scores groups together the knockdown, the parent, and the rescue cells (Fig. 5B). Most of the methylation sites affected by Nopp140 KD (red in the dendrogram) cluster with the hypomodified sites in rRNA (Fig. 5B).

**Figure 5.**
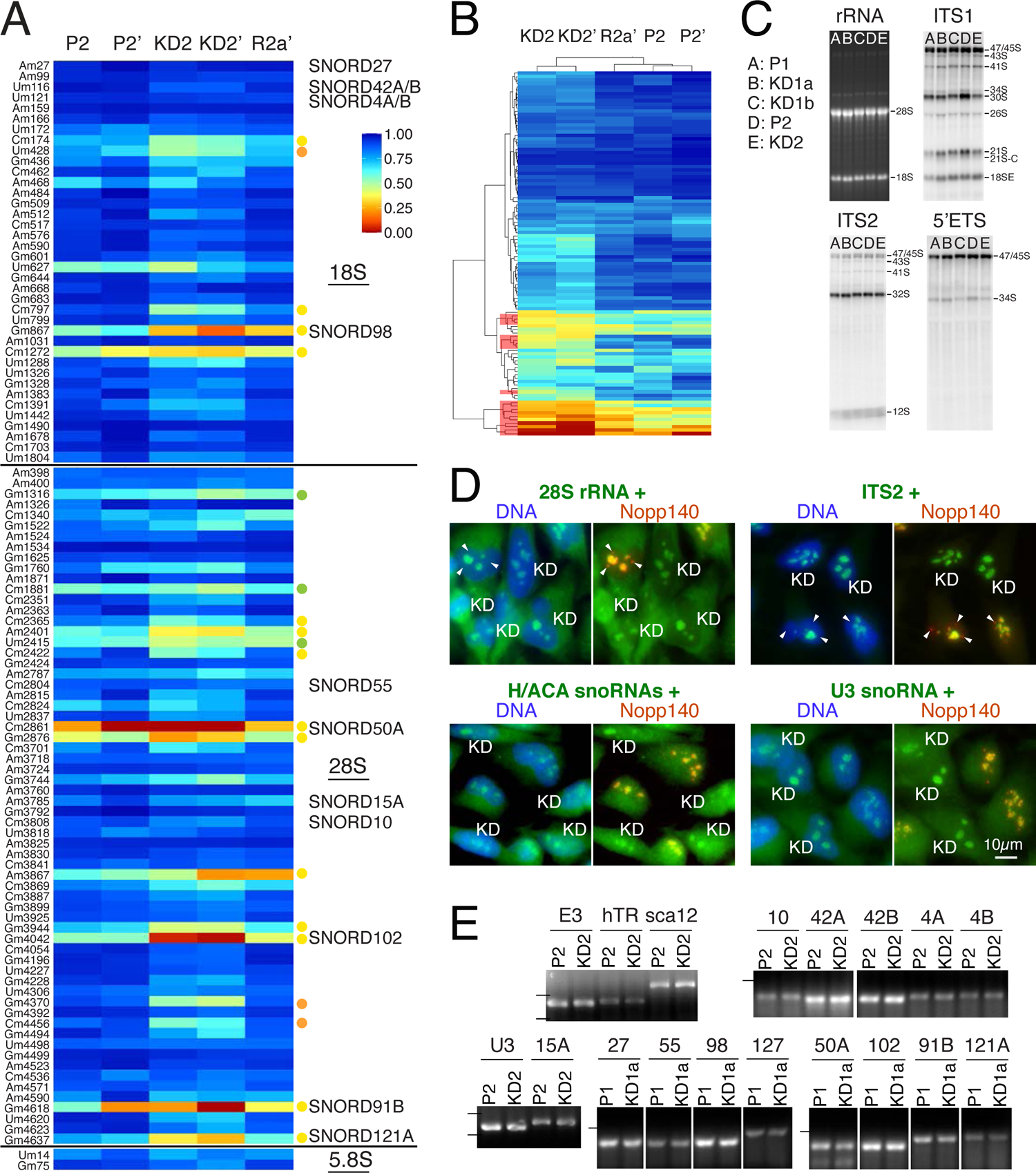
RiboMethSeq (RMS) of rRNAs shows reduced 2’-O-methylation at select sites without consequences for pre-rRNA processing, localization, or steady-state levels of snoRNAs and mature rRNAs between parent and KD cells. (A) Heatmap of RMS scores from two experiments as in Fig. 4A but of rRNAs. Sites of reduced 2’-O-methylation are indicated with yellow or, if normally fully modified, with orange dots. Normally hypomethylated nucleotides that are unaffected by Nopp140 KD are indicated with green dots following the color scheme of Supplemental Table S2. The snoRNAs tested for their abundance in (E) are indicated by name next to the nucleotide they are complementary to for site-directed modification. (B) Unsupervised clustering of the values in (A) groups the KD cells together and shows the sites of reduced 2’-O-methylation (red in the dendrogram on the left) to segregate with the normally hypomodified residues in the parent and rescue cells. (C) Mature rRNA and pre-rRNA processing analysis. Total RNA (3 µg) from the indicated cell lines was separated on denaturing agarose gels, stained with ethidium bromide, to detect mature 18S and 28S rRNAs, or processed for northern blotting with specific probes (ITS1, ITS2, and 5’ ETS), to detect all major precursor rRNAs. Note, the data shows no significant differences in mature rRNA production or pre-rRNA processing between the parent and KD cells. (D) RNA fluorescence in situ hybridization (FISH; green) combined with indirect immunofluorescence of Nopp140 (red/yellow; right panels) and DAPI DNA stain (blue; left panels) of a 1:1 mixture of parent and KD cells grown side by side. The detected RNAs are indicated on top of each pair of panels. Parent cells are recognized in the right panels by the nucleolar staining of Nopp140 (yellow because or overlap with green RNAs) and the knockdown cells are labeled (KD). Some CBs are indicated (arrowheads) and appear in red in the Nopp140 stain because they are devoid of (pre)-rRNAs. A combination of P1 and KD1a cells (28S rRNA and U3 snoRNA) and of P2 and KD2 cells (ITS2 and H/ACA snoRNAs) were used. To detect H/ACA snoRNAs, a combination of primers was used against E3, ACA8, ACA18, and ACA25. Scale bar = 10µm. (E) Analysis of steady-state levels of select C/D and H/ACA RNAs by semi quantitative RT-PCR. RNAs tested and cell pairs analyzed are indicated above each lane. A simple number refers to that specific SNORD. The dashes on the left indicate the migration position of markers, 100nts if only one and 100 and 200nts if two.

To assess a possible impact on ribosome biogenesis of these subtle but reproducible changes in rRNA 2’-O-methylation, we investigated pre-rRNA processing by northern blotting with probes for specific processing intermediates on total RNA from parent and KD cells (Fig. 5C).

Specifically, we employed probes for the internal transcribed spacer 1 (ITS1) and 2 (ITS2) and for the 5’-external transcribed spacer (5’ ETS) (Fig. 5C). No significant variations in mature 28S and 18S rRNAs and in pre-rRNA processing were detected between the parent and KD cell lines (Fig. 5C). We further examined if the changes in rRNA 2’-O-methylation impacted the localization of the rRNAs themselves or of snoRNAs. For this purpose, we grew parent (Nopp140-positive) and KD cells (Nopp140-negative) on the same dish in a 1:1 mixture (Fig. 5D). We detected 28S rRNA, ITS2, H/ACA snoRNAs (E3, ACA8, ACA18, and ACA28 combined), and the C/D snoRNA U3 by RNA FISH (Fig. 5D, both panels, green) together with immunofluorescence for Nopp140 (right panels, red) to identify parent (orange nucleoli) and KD cells (green nucleoli, KD). Note the localization of 28S rRNA in both its place of synthesis (nucleoli) and of function (cytoplasmic ribosomes) (Fig. 5D). In contrast, ITS2 was restricted to nucleoli, its place of synthesis and processing. As expected, neither 28S nor ITS2 rRNAs were detected in CBs (Fig. 5D, arrow heads in Nopp140-positive cells). Localization of none of the RNAs changed between parent and KD cells (Fig. 5D). The absence of any difference in these assays was perhaps unsurprising considering the absence of any notable pre-rRNA processing and growth defects in Nopp140 KD cells.

To investigate the mechanism underlying the reduction in rRNA methylation at only a few residues, we used semi-quantitative RT-PCR to interrogate the levels of some of the snoRNAs responsible for specifying the modifications. For this purpose, we isolated total RNA from parent and Nopp140 KD cells and used sno/scaRNA-specific primers in RT-PCR reactions. As previously established, scaRNAs displaced from CBs did not change in their abundance, e.g., hTR and scaRNA12/U89 (Bizarro et al., 2019) nor did that of the H/ACA snoRNA E3/SNORA63 (Fig. 5E). Similarly, 14 nucleolar box C/D snoRNAs did not vary in abundance between parent and KD cells irrespective if 2’-O-methylation of their target residue varied (SNORD98, 127, 50A, 102, 91B, and 121A) or not (SNORD10, 42A, 42B, 4A, 4B, 15A, 27, and 55) (Fig. 5E). This despite the fact that for some targets, methylation was reduced by over 50% (SNORD50A, 102, 91B, and 121A) (Supplemental Table S2). Additionally, the levels of the box C/D snoRNA U3, which is involved in pre-rRNA processing, did not vary between parent and KD cells (Fig.5E) nor did its localization (Fig. 5D). The results were independent of which pairs of cell lines were compared, P2 vs. KD2 or P1 vs. KD1a (Fig. 5E). Consequently, the reduced levels of 2’-O-methylation of certain nucleotides is not caused by fluctuations in levels of the snoRNPs that guide their modifications. This absence of correlation between snoRNA abundance and methylation levels observed here is consistent with previous studies employing RMS (Krogh et al., 2016; Sharma et al., 2017). However, the low levels of Nopp140 in nucleoli leading to decompaction of the DFC apparently modified the access of snoRNPs to their targets affecting particularly those responsible for the methylation of normally already hypomodified residues. This may be similar to Nopp140 corralling those scaRNPs in CBs whose function there is required for snRNA modification, as compared to those that can also function in the nucleoplasm.

### Effects of Nopp140 KD on mRNA expression and pre-mRNA splicing fidelity

The impact of Nopp140 KD on specific sites of 2’-O-methylation in several spliceosomal snRNAs provided us with the opportunity to test for the first time the function of these modifications in pre-mRNA splicing. For this purpose, total RNAseq was performed on all cell lines. The reproducibility and robustness of our sequencing data was remarkable (e.g., Fig. 6E and 7B), given that total RNA from every cell line was isolated three times and as sequencing of the P2 and KD2 and of the P1, KD1a, and KD1b cell lines was performed one year apart and on different continents.

**Figure 6.**
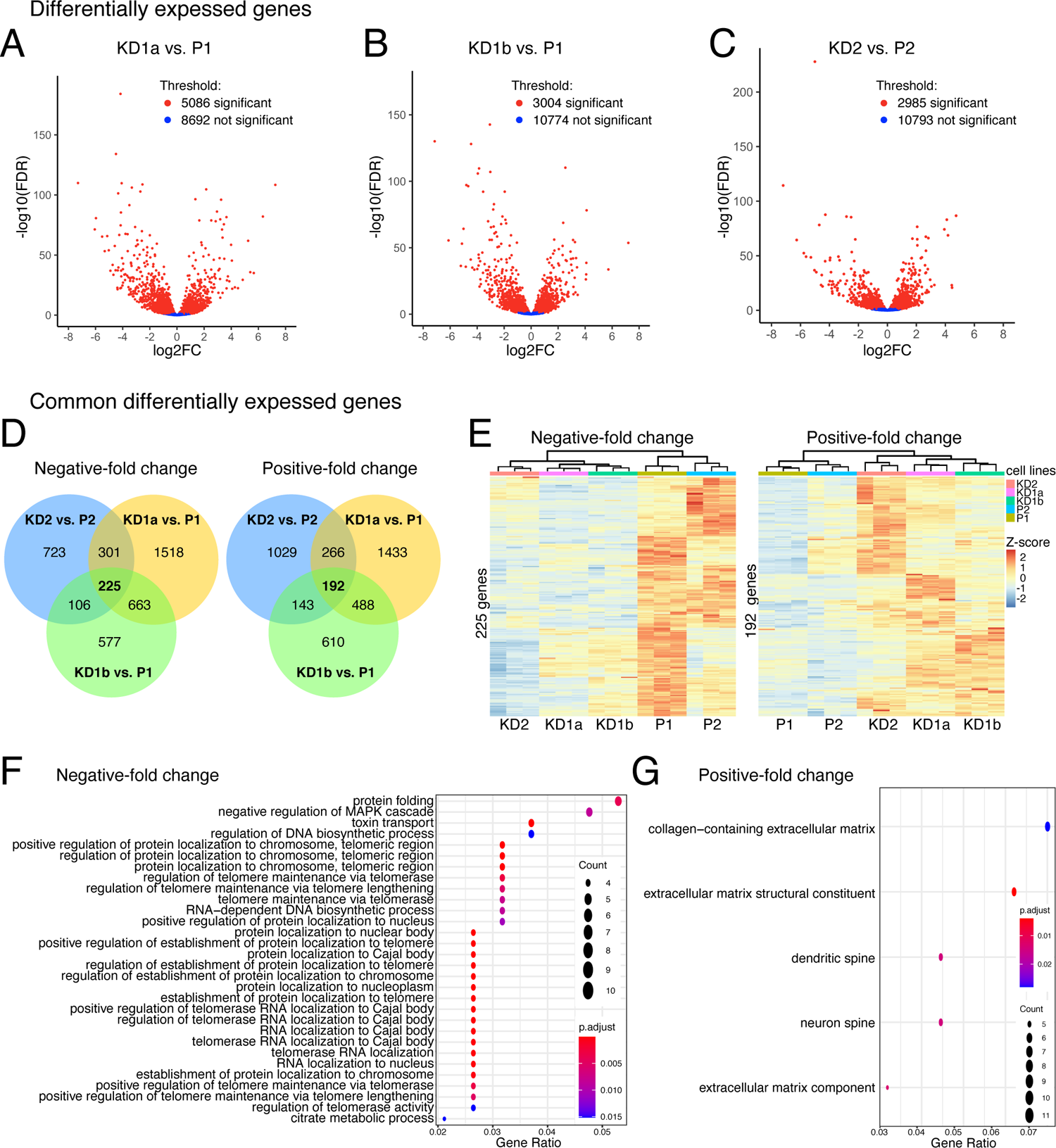
RNAseq of total RNAs from Nopp140 parent and knockdown (KD) cells reveals few common differentially expressed genes. (A) Volcano plots of genes differentially expressed between KD1a and P1 cells, (B) between KD1b and P1 cells, and (C) between KD2 and P2 cells. (D) Venn diagrams for common differentially expressed genes between the 3 pairs in (A) to (C) for negative- and positive-fold change. (E) Heatmap of z-scores for 225 common genes with negative- and 192 common genes with positive-fold change for all 5 cell lines. Note, despite the small differences, all biological triplicates and the parents and KD cells cluster together, even those KD cells originating from the same parent, KD1a and b. (F) Gene ontology (GO) analysis of common differentially expressed genes with a negative-fold change reveal mostly genes related to Nopp140 function. (G) GO analysis of common differentially expressed genes with a positive-fold change reveals only few genes. Gene ratio is the percentage of total differentially expressed genes within a given GO term.

**Figure 7.**
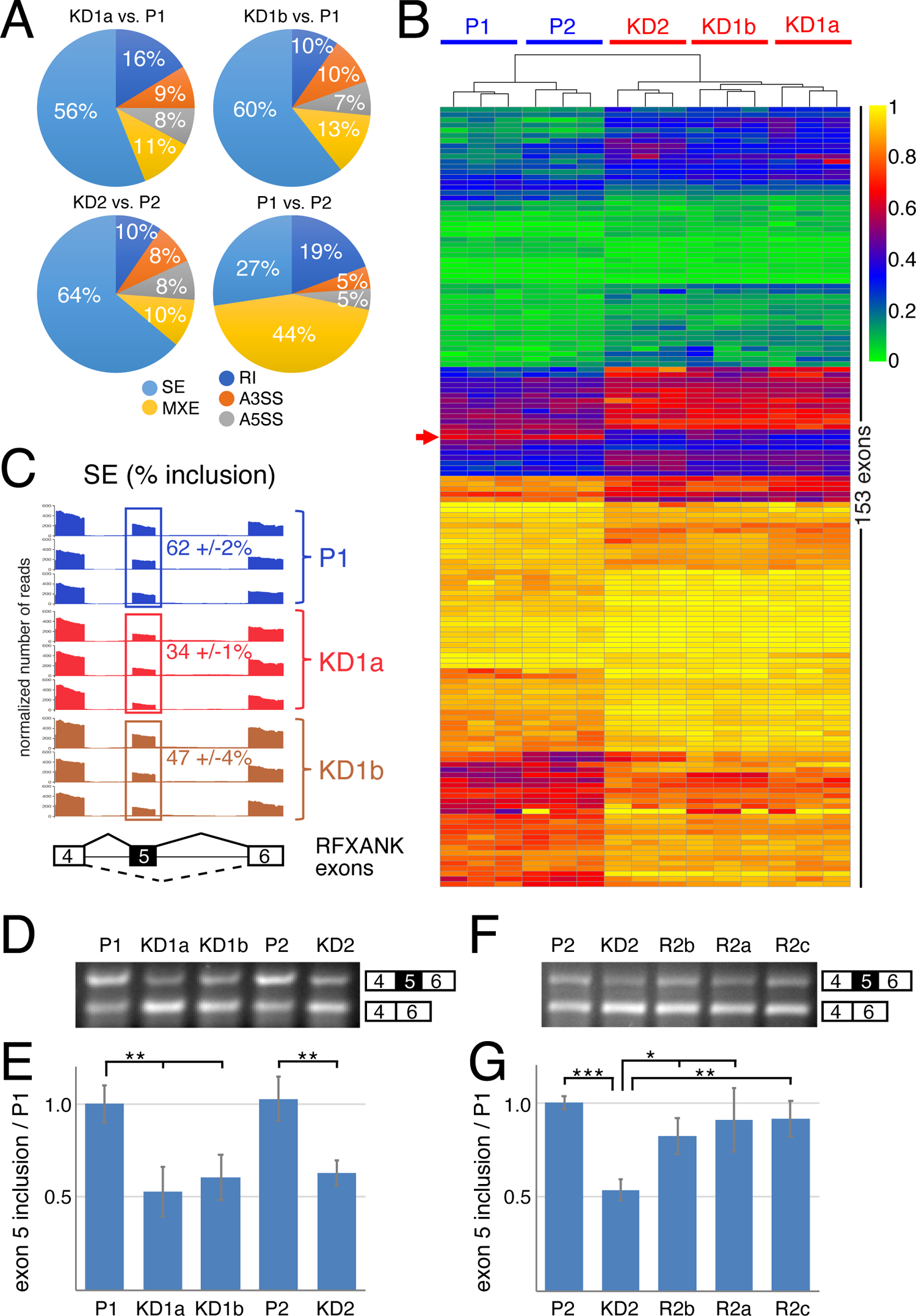
Analysis of RNAseq data for alternative splicing events using the rMATS algorithm shows small but significant changes after Nopp140 KD. (A) Analysis of splicing events of Nopp140 KD cells relative to their corresponding parent cells contrasted to those normally occurring in parent versus parent cells. The analyzed events expressed in the percentage pie chart are skipped exons (SE), mutually exclusive exons (MXE), retained introns (RI), alternative 3’-splicesite (A3SS), and alternative 5’-splicesite (A5SS). (B) Heatmap of 153 alternatively spliced exons in all cell lines arranged through unsupervised clustering. Note the remarkable clustering of all triplicates and the parent versus KD cells. The RFXANK exon 5 is pointed out (red arrow). (C) Sashimi plot of the sequence traces spanning the region of RFXANK exons 4 – 6 (schematically depicted underneath) from triplicate analysis. For direct comparison, the reads are normalized and the mean inclusion levels (+/- SD) of exon 5 (boxed) are indicated. Note the larger difference of exon 5 inclusion of KD1a over KD1b cells relative to the parent P1 mirroring the degree of Nopp140 KD in those cells. Also note the small SD values. The plots were generated using ggsashimi (https://github.com/guigolab/ggsashimi). (D) Semi-quantitative RT-PCR of the RFXANK exons 4 – 6 separated on agarose gels and stained by ethidium bromide confirm the deep sequencing rMATS results for exon 5 inclusion. (E) Quantification of quadruplicate results in (D). Unpaired t-tests identify significant differences (*** P < 0.005). (F) As in (D) but including RNA from all 3 rescue cell lines documenting the restoration of exon 5 inclusion in Nopp140 re-expressing cell lines. (G) Quantification of triplicate results in (F). Unpaired t-tests identify significant differences (* P < 0.05, ** P < 0.005, *** P < 0.0005).

We first investigated differential expression of genes in the KD cells versus their parent cells. Whereas the number of up and downregulated genes was similar for each of the three KD and parent pairs, the overall number of significantly differentially expressed genes (FDR 0.05) between KD1a and P1 was 5086 and that for KD1b and P1 was 3004 out of 13,778 genes analyzed (Fig. 6A and B). Similar numbers of differentially expressed genes were noted between P2 and KD2 cells (Fig. 6C). Limiting our analysis of differentially expressed genes to those common between all three pairs yielded 225 downregulated and 192 upregulated genes (Fig. 6D). Unsupervised clustering of these genes grouped together each triplicate RNAseq, the parents P1 and P2, the KD cells KD1a, KD1b, and KD2 (Fig. 6E). Even the KD cells derived from the same parent cells, KD1a and KD1b, grouped together (Fig. 6E). Interrogating the differentially expressed genes according to gene ontology revealed almost exclusively genes associated with Nopp140 function. Specifically, downregulated genes were highly enriched in genes involved in localization to CBs, nuclear bodies, nucleus, chromosomes, and telomeres (Fig. 6F). Apparently, in the absence of Nopp140, there is a reduced need for these co-depleted genes. For unknown reasons, upregulated genes showed a slight enrichment in genes involved in extracellular matrix and neuronal spine formation (Fig. 6G).

To assess the impact on alternative splicing of Nopp140 KD, i.e., the loss of 2’-O-methylation from several spliceosomal snRNAs, we employed the rMATS algorithm for analysis of our RNAseq data. Of the 5 different alternative splicing events, skipped exon (SE), retained intron (RI), alternative 3’ and 5’ splice sites (A3SS and A5SS), and mutually exclusive exons (MXE), SE events far outnumbered the others with over 50% in comparisons of all three parent and KD sets (Fig. 7A). This was in stark contrast to a comparison of the two parent cells where mutually exclusive exons with 44% far outnumbered skipped exons and the other splicing events (Fig. 7A). Therefore, reduced 2’-O-methylation of these snRNA residues preferentially affected one splicing pathway. However, only 153 SE events were common to all three sets indicating that general splicing remains unaltered (Fig. 7B). When analyzed by unsupervised clustering, each triplicate sample, the two parents, and the three KDs, all grouped together again highlighting the robustness of the data (Fig. 7B). Gene ontology analysis of the genes harboring the 153 SE events, did not reveal any significantly enriched GO terms. For further analysis and corroboration of the data, we focused on the RFXANK gene whose exon 5 was skipped about twice as often in the KD1a cells compared to the parent. Specifically, exon 5 was included in 62 ± 2% of mRNAs in parent P1 cells as shown on a sashimi plot (Fig. 7C). In contrast, the Nopp140 KD1a and KD1b cells included exon 5 only in 34 +/-1% and 47 +/-4% of mRNAs, respectively (Fig. 7C). Thus, also the degree of exon skipping parallels that of Nopp140 KD (Fig. 1A and B) and that of the loss of snRNA 2’-O-methylation in those cells (Fig. 3-5). In other words, exon skipping is more pronounced in KD1a than in KD1b cells (Fig. 7C) suggesting that these reproducible changes in alternative splicing are a direct consequence of reduced snRNA 2’-O-methylation. We corroborated the RNAseq data by semiquantitative RT-PCR (Fig. 7D).

Quantification of the results expressed relative to P1 cells showed a loss of exon inclusion by about 50% (Fig. 7E) mirroring the RNAseq results (Fig. 7C). Importantly, exon 5 inclusion was restored in all 3 rescue cell lines (Fig. 7F and G), which paralleled the rescue of 2’-O-methylation of U1, U2, U5, and U12 snRNAs (Fig. 4C and Supplemental Table S1). These data thus strongly support the importance of snRNA modification for maintaining the fidelity of pre-mRNA splicing.

## DISCUSSION

In this study, we took advantage of our ability to separate the bulk of scaRNPs from their target snRNAs in CBs. This was achieved by KD of Nopp140 or expression of unphosphorylated Nopp140, resulting in CBs devoid of Nopp140 and scaRNPs. This approach allowed us to study the function of CBs in snRNA modification. We observed that in Nopp140 KD cells, snRNA pseudouridylation proceeds normally but most sites of 2’-O-methylation were impaired. Genome wide analysis shows an overall switch in pre-mRNA splicing events from mutual exclusion of exons to exon skipping and a reproducible change in alternate splice site usage. All these effects are due to Nopp140 KD as their degree parallels that of KD levels and because Nopp140 re-expression rescues all effects. Two main conclusions can be drawn from these observations, most snRNA 2’-O-methylation occurs in CBs and is required to maintain splicing fidelity.

A surprising finding was the specific effect on 2’-O-methylation but not on pseudouridylation, at least not at the interrogated sites. These data support the notion that CBs are not obligatory sites for pseudouridylation of snRNAs but that this modification can also occur in the nucleoplasm. In fact, in Drosophila and HeLa cells, snRNAs are efficiently modified even in the absence of CBs (Deryusheva and Gall, 2009, 2013; Deryusheva et al., 2012). In our cell system, coilin-positive CBs with snRNPs persist even without scaRNPs and Nopp140 (Bizarro et al., 2019). The modifications still occurring, therefore, appear essential for pre-mRNA splicing. In particular, pseudouridines in U2 snRNA have long been identified as important for in vitro reconstituted splicing reaction in yeast extracts (McPheeters et al., 1989). Pseudouridines of yeast U2 snRNA are further important for stimulating the ATPase activity of Prp5 during spliceosome assembly (Wu et al., 2016). Finally, pseudouridines in the branch site recognition region of U2 snRNA and in general are required for pre-mRNA splicing and snRNP biogenesis in vivo and in vitro (Yu et al., 1998; Zhao and Yu, 2004, 2007).

In contrast, most 2’-O-methyl groups of snRNAs U1, U2, U5, and U12 are lost in Nopp140 KD cells allowing investigation of their role in pre-mRNA splicing. As Nopp140 KD cells proliferate at identical rates as their parent cells (Bizarro et al., 2019), 2’-O-methylations of snRNAs appear less important than pseudouridines for overall splicing. This is supported by the fact that in yeast, snRNAs are pseudouridylated, but 2’-O-methylation has not been observed (Massenet et al., 1998). Nevertheless, we show that reduced 2’-O-methylation of snRNAs compromises alternative splicing. Apparently, 2’-O-methylation in the nucleoplasm is not as efficient in the absence of CBs arguing that CBs function by enhancing the activity of scaRNPs on snRNAs by bringing them together following the law of mass action. Apparently, an opposite function of CBs is in place for the telomerase RNP, in which case CBs function to sequester telomerase from its nucleoplasmic substrates, the telomeres (Bizarro et al., 2019)

Interestingly, two 2’-O-methyl groups within U2 snRNA are not impacted by Nopp140 KD, Gm12 and Gm25. In the active spliceosome, those 2’-O-methylated residues lie between helix Ia and Ib, and right adjacent to helix II formed between U2 and U6 snRNAs, perhaps pointing to an especially important role in splicing (Townsend et al., 2020; Zhang et al., 2017). In fact, U2-Gm12 was one of four 2’-O-methylated residues within the first 20 nucleotides of HeLa U2-snRNA that was required for in vitro splicing (Dönmez et al., 2004). U2-Gm12 and U2-Gm25 form the base of stem loop I (SL1) and of the branchpoint-interacting stem loop (BSL), respectively, in the 17S U2 snRNP (Zhang et al., 2020). However, they are both located in the center of SL1 of the extended U2 conformation in the active spliceosome. Apparently, the modifications of these two guanosines exhibit a more basic function in splicing than the remaining ones of U2 and of all other snRNAs. Similarly, pseudouridines cannot be lost from snRNAs, although to determine if this is true for all pseudouridines will require a more detailed genome-wide approach, perhaps using the recently developed HydraPsiSeq (Marchand et al., 2020).

Methylation of 2’-O-ribose of U6 snRNA remained unaffected in the Nopp140 KD cells. This is not surprising given that its modification occurs in nucleoli and not CBs (Ganot et al., 1999). In fact, guiding the 2’-O-methylation of U6 snRNA follows an altogether different pathway than that for all other snRNAs. The La related protein 7 (LARP7) is responsible for bringing together U6 snRNA and a specific subset of C/D snoRNAs required for its modification (Hasler et al., 2020). Thus, LARP7 may function for U6 in the nucleoplasm as Nopp140 does for all other snRNAs in CBs.

While overall 2’-O-methylation of snRNAs is clearly reduced in Nopp140 KD cells that of rRNA is affected to a much lesser extent. Only 13% and 18% of 2’-O-methylated residues of 18S and 28S rRNA, respectively, are impacted by Nopp140 KD. This is in stark contrast to yeast cells, where significant reduction of 2’-O-methylation at about half of all normally modified rRNA sites was observed after depletion of the helicases Prp43 or Dbp3 (Aquino et al., 2021). Importantly, all but three of the sites affected by Nopp140 KD are normally not fully 2’-O-methylated in cellular rRNA. Apparently, these hypomodified residues are generally less important for proper ribosome biogenesis and function and thus more susceptible to minor changes in their cellular environs. This conclusion seems similar to the detrimental effect on general pre-mRNA splicing by the loss of snRNA pseudouridylation but not the reduction of snRNA 2’-O-methylation.

Regardless, as in the case of the two guanosines of U2 snRNA that fail to lose their 2’-O-methyl groups, the loss of those three nucleotides normally modified to the full extent may be the ones to look at for consequences. Given that the Nopp140 KD cells exhibit the same growth rate as their parent cells, it is not surprising that no obvious function can be assigned to those three nucleotides, except that they seem to be positioned near the subunit interface.

The low impact on rRNA but high impact on snRNA modification by Nopp140 KD may be further explained by the presence of residual Nopp140 in nucleoli but not CBs. This was particularly surprising given that in parent cells Nopp140 fluorescence intensities of CBs matched those of nucleolar DFCs indicating a specific loss of Nopp140 from CBs but only a reduction in nucleoli. The extreme transcription rate of rDNA and the concomitant near normal rRNA modification in regularly growing Nopp140 KD cells draws a typical accumulation of snoRNPs in DFCs, which may be responsible for the visible Nopp140 accumulation even in KD cells. In contrast, as outlined above, most scaRNPs are displaced from CBs into the nucleoplasm in KD cells where residual Nopp140 is more dispersed and less detectable. In other words, nucleoli are obligate cellular organelles whereas CBs are not but may simply enhance activity by co-concentration of scaRNP enzymes and their snRNA substrates. This is supported by ample data of cells functioning and splicing normally even in the absence of CBs (Deryusheva and Gall, 2009, 2013; Deryusheva et al., 2012; Spector et al., 1992).

The robustness of the overall RNAseq results is remarkable and with it that of pre-mRNA splicing analysis using the rMATS algorithm. In particular, unsupervised clustering not only aligns the biological triplicates of each cell line but also clusters the parent and KD cells together, even the KD cells derived from the same parent cells. This despite the fact that library preparation and sequencing of the two parent–KD cell pairs was performed a year apart and on different continents. Although overall splicing was minimally affected, there were some obvious differences between parent and KD cells. There was a clear shift in preference for splicing events when parent cells were compared to KD versus each other. In particular, comparison to KD cells showed a strong preference for exon skipping events. Overall, 153 alternatively spliced exons were identified consistently in all three KD cells highlighting the effect of reduced snRNA 2’-O-methylation on pre-mRNA splicing. Importantly, the degree of effect paralleled that of reduction in snRNA 2’-O-methylation and that of Nopp140 KD levels clearly linking these events. This was further corroborated by the rescue of all effects by re-expression of Nopp140. Mechanistically, the entire chain of events depends on the Nopp140-mediated concentration of scaRNPs (and Nopp140) in CBs, which we show relies on the extreme level of phosphorylation of Nopp140.

Nopp140 phosphorylation at ∼80 serines is mediated by CK2 and is required for accumulation in CBs, but not nucleoli. Apparently, phosphorylation of Nopp140 is not sufficient for its interactions with snoRNPs in nucleoli but is for those with scaRNPs in CBs. Therefore, inhibition of Nopp140 phosphorylation specifically inhibits scaRNP localization to CBs and with it most 2’-O-methylation of snRNAs resulting in altered splicing fidelity. These phosphorylation specific effects seem surprising given that normally dephosphorylation of Nopp140 is not observed and consequently only newly synthesized Nopp140 appears a target for CK2. Nevertheless, these effects need to be taken into consideration in therapy with the CK2 inhibitor CX-4945 (silmitasertib) for cholangiocarcinoma, hematological malignancies, and COVID-19.

## MATERIALS AND METHODS

### Plasmids

Plasmids used to generate the stable rescue clones are pNK65 and pJB9 expressing HA-Nopp140-GFP under a CMV promoter or UBC promoter, respectively. Transient rescue during CKII inhibition was performed using plasmid pJB8 expressing HA-Nopp140 under CMV promoter (Bizarro et al., 2019).

### Cell culture, transfection and genome engineering

HeLa cells and the various clones were cultured in DMEM (Gibco) and 10% fetal bovine serum (Atlanta Biologicals) at 37°C under 5% CO2 in air. Nopp140 KD clones were generated as described using CRISPR/Cas9 technology (Bizarro et al., 2019). RNAseq in this study confirmed proper targeting of the sgRNAs with most reads carrying mutations at the targeted sites. Transfections were performed with Lipofectamine 3000 (Thermofisher Scientific) following the manufacturer’s protocol. CK2 inhibition was performed on HeLa cells or on Nopp140 KD1a cells transfected with pJB8 plasmid six hours prior to addition of CX-4945 (Selleckchem) at 10µM in DMEM, 10% FBS for one, two or three days before analysis by western blotting and indirect immunofluorescence. Stable rescues were generated in the Nopp140 KD2 cells by transfection with pJB9 or pNK65 plasmids. Transfected cells were treated with G418 (1g/ml final) (Corning) for 2 months. Single clones were obtained by limited dilution and tested for Nopp140 re-expression by indirect immunofluorescence and western blotting. Transfection with pJB9 yielded clones R2a and R2b and that with pNK65 yielded clone R2c. Nopp140 expression in the rescue clones remained stable after several months in culture.

### Antibodies

Antibodies (dilutions in parentheses) for western blotting (WB) or indirect immunofluorescence (IF) were as follows: anti-Nopp140 rabbit serum (RS8 at 1:5000 for WB and 1:1000 for IF) (Kittur et al., 2007); mouse monoclonal anti-NAP57 immunoglobulin G (IgG) (H3 at 1:500 for IF; Santa Cruz Biotechnology); mouse monoclonal anti-β-actin (AC-15 at 1:1000 for WB; Santa Cruz Biotechnologies); mouse anti–γ-tubulin ascites fluid (GTU-88 at 1:5000 for WB; Sigma); DyLight488 goat anti-mouse IgG (1:500 for IF) and rhodamine (TRITC) goat anti-rabbit IgG (1:500 for IF; both Jackson Immuno Research); Alexa Fluor 680 goat anti-rabbit IgG (1:10,000 for WB; Thermo Fisher Scientific); IRDyeTM 800 goat anti-mouse IgG (1:10,000 for WB; Rockland Immunochemicals).

### Western blotting

For each experiment, proteins from the same number of cells per condition were extracted into SDS-sample buffer (0.5 M Tris; pH6.8; 12% SDS; 0.05% bromophenol blue). The lysates were tip sonicated and total proteins loaded (100,000 cell equivalents), separated on 9% SDS– PAGE, and transferred to nitrocellulose membrane. Transfer efficiency was confirmed by Ponceau red staining, and membranes were blocked in blocking buffer (Tris-buffered saline, 0.1% Tween, and 2.5% nonfat dry milk) for 30 min before incubation with primary antibodies diluted in blocking buffer overnight at 4°C. After 3 washes in blocking buffer, membranes were incubated with appropriate secondary antibodies diluted in blocking buffer for 1 h at room temperature in the dark. After 3 washes in blocking buffer, membranes were scanned on an Odyssey 9120 Imaging System (LI-COR Biosciences), and protein bands were quantified using Image Studio Lite (LI-COR Biosciences) and analyzed with Microsoft Excel and GraphPad Prism software.

### Indirect immunofluorescence

Cells grown on coverslips were fixed in 4% paraformaldehyde (PFA) in phosphate-buffered saline (PBS) for 20 min, permeabilized with 1% Triton X-100 in PBS for 5 min, and blocked with 1% powdered milk in PBS (IF blocking buffer) for 15 min. The cells were then incubated for 2 h with primary antibodies in IF blocking buffer, washed, and incubated for 1 h with secondary antibodies in IF blocking buffer in the dark. This was followed by washing and DNA staining with 4’,6-diamidino-2-phenylidone (DAPI; 1 µg/ml in PBS). Coverslips were mounted on glass slides using ProLong Diamond Antifade Mount (Thermo Fisher Scientific) and observed using a Zeiss Axio Observer Z1 fluorescence microscope (63x objective, NA 1.4) with filter sets 34-DAPI (Zeiss #000000-1031-334), 10-AF488 (Zeiss #488010-9901-000), 43HE-DsRED (Zeiss #489043-9901-000), and 50-Cy5 (Zeiss #488050-9901-000). Z-stack images in 200-nm steps were acquired with a Zeiss AxioCam MRm camera using Axiovision software (Zeiss). Maximum projections were generated using ImageJ (National Institutes of Health). Quantification of Nopp140 protein signals in nucleoli and Cajal bodies was done using ImageJ with the help of macros (available upon request). Briefly, NAP57 images were used to locate the nucleoli and Cajal bodies around which masks were generated. The DAPI images were used to establish nuclear masks. These masks were applied to the Nopp140 images to determine their signal intensity in the organelles per nucleus. Background was subtracted individually for each nucleolus and Cajal body and was defined as the pixel with the lowest signal in a 50-pixel circumference of the mask using an ImageJ function. Images for figures were cropped and adjusted using Photoshop CC (Adobe). To compare parent, KD, and rescue cell images, all images within the same panels and of the same antigens were acquired and adjusted identically.

### RNA fluorescent in situ hybridization (FISH)

Cells on coverslips were fixed with PBS, 4% PFA, permeabilized with PBS, 1% Triton-X100, washed with 2xSSC, 40% formamide. RNAs were stained for 4 hours at 37°C in the dark using 32-50mer probes (Supplemental Table S3) synthesized and internally labeled with Cy3 as described (Chartrand et al., 2000). Hybridization with denatured probes (2ng/µL) was performed for 3 hours at 37°C in 2xSSC, 40% formamide, 50ng/µL ssDNA/tRNA, 3.5µg/µL BSA. Cells were washed extensively with 2xSSC, 40% formamide at 37°C, then with PBS, before fixation in PBS, 4% PFA and incubation in blocking buffer (PBS, 1% dry milk). Cells were then incubated with Nopp140 antibodies (RS8 at 1:1000) in blocking buffer for 2 h followed by secondary antibodies (Rabbit-Alexa488 at 1:500) in blocking buffer for 1 h in the dark. After washes in blocking buffer and DAPI staining, the coverslips were mounted using ProLong Diamond. The samples were observed using an Olympus IX81 epifluorescence microscope with Objective 60x, 1.4NA, oil-immersion objective. Z-stack images in 200-nm steps were acquired with a Sensicam QE cooled CCD camera using IP Lab 4.0.8 software and processed using Adobe Photoshop.

### Total RNA extraction and sequencing

RNA from the different cell lines was extracted using 500 µl TRIzol reagent (Ambion) directly on 10-cm dishes (cell confluency ∼80%, ∼1,000,000 cells). Lysed cells in TRIzol were scraped into tubes, extracted twice with chloroform, the RNA was precipitated with 0.7 volume isopropanol after addition of 20 µg glycogen, and resuspended in UltraPure distilled water. RNA concentration and quality were determined by Nanodrop (ratio 260/230 and 260/280 above 1.8) and Agilent 2100 BioAnalyzer (RIN above 8). Total RNA was used for northern blot analysis, RiboMethSeq, RT-PCR, and RNase H analysis. For deep sequencing, total RNA was prepared from 3 separate dishes for each sample and shipped to Novogene Corporation Inc. (Sacramento, CA) for cDNA library preparation (250∼300 bp inserts) and Illumina sequencing (PE150). The RNA was prepared and sequenced one year apart in the USA and in China in two batches, one for the P1, KD1a, and KD1b cells and one for the P2 and KD2 cells. All data is deposited in the GEO repository under the accession number GSE173171.

### RNAseq data analysis

Raw Fastq files were obtained from NOVOGENE and were checked for quality of reads with FastQC (version 0.11.4). Raw reads were aligned with the splice aware aligner STAR (version 2.4.2a) (Dobin et al., 2013). Cufflinks (version 2.2.1) was used to generate FPKM expression values (Trapnell et al., 2010). The featureCounts from Subread package (1.5.0-p1) was used to count the number of raw fragments associated with each gene (Liao et al., 2014). Differential gene expression analysis was performed with the help of the Bioconductor package DESeq2 (Love et al., 2014). Gene ontology (GO) analysis of common differentially expressed genes was performed using clusterProfiler (Yu et al., 2012). Multivariate Analysis of Transcript Splicing with replicates (rMATS, version 3.2.5) was used to detect differential splicing events. Significant events with FDR ≤ 0.05 are reported (Shen et al., 2014).

### Pre-rRNA processing analysis by Northern blotting

Total RNA was extracted with the TRIzol reagent (Ambion) according to the manufacturer’ instructions. 3 µg total RNA was separated on 1.2% denaturing agarose gel, transferred to a nylon membrane, and hybridized with 32P-labelled oligonucleotide probes specific to all major pre-rRNAs, as described in (Sharma et al., 2015; Tafforeau et al., 2013). The membrane was exposed to Fuji imaging plates (Fujifilm). The signals were acquired with a Phosphorimager (FLA-7000, Fujifilm) and quantified with the native Multi Gauge software. Probe sequences are provided in Supplemental Table S4.

### Analysis of 2’-O-methylation levels by RiboMethSeq

RMS was performed exactly as described in (Marchand et al., 2016). 150 ng total RNA was used in each reaction. The samples were sequenced at the ULB-BRIGHTcore facility (Brussels Interuniversity genomics high throughput core) on Illumina Novaseq 6000 as paired-end runs (100 nt read length). In average 25 million reads were sequenced (Supplemental Table S5).

Adapter sequences were removed using Trimmomatic (0.36; LEADING:30 TRAILING:30 SLIDINGWINDOW:4:15 MINLEN:17 AVGQUAL:30) and reads in forward direction were mapped to an artificial genome containing ribosomal RNA sequences using bowtie2 (2.3.3.1; sensitive). Mapped reads were analyzed using the R package version 1.2.0 “RNAmodR.RiboMethSeq: Detection of 2’-O-methylations by RiboMethSeq” (Ernst F.G.M. and Lafontaine D.J.L. 2019) and the Score C/RiboMethScore was used as a measurement for 2’-O methylation. For the analysis of methylation levels on positions known to be methylated, data from the snoRNAdb was used (Lestrade and Weber, 2006) and updated to revised rRNA sequence coordinates based on NCBI accession NR_046235.3, which are available from the ‘EpiTxDb’ R package version 1.0.0 (Ernst F.G.M. 2020).

### Fluorescent primer extension analysis of RNA modification

To analyze 2’-O-methylation and pseudouridylation patterns of human U2 snRNA we used a fluorescent primer extension method as described (Deryusheva and Gall, 2009; Deryusheva et al., 2012). In brief, to detect 2’-O-methylated positions reverse transcription reactions were performed at very low concentration of dNTP. It has been shown previously that different reverse transcription enzymes have different termination efficiency at 2’-O-methylated positions (Deryusheva et al., 2012). We used EpiScript RT (Epicentre) to assess U2-Am30 semi-quantitatively, and AMV-RT (New England Biolabs) to assess U2-Um47. To map pseudouridines, RNA samples were treated with CMC (N-cyclohexyl-N9-[2-morpholinoethyl] carbodiimide metho-p-toluene sulfonate) followed by incubation in sodium carbonate buffer (pH 10.4). Reverse transcription was done at 0.5 mM dNTP using EpiScript RT.

Each RNA sample was tested in 2-3 replicates. Sets of RNAs from parental and Nopp140KD lines were always treated simultaneously and separated on capillary columns in parallel using serial dilutions. GeneMapper 5 software (Applied Biosystems) was used to visualize and analyze the data.

### RT-PCR-based RNA modification analysis

Another RT-based method to assay RNA modification levels semi-quantitatively utilizes the same principles as described above but instead of separation of fluorescently labeled ssDNA fragments on capillary columns it involves qPCR with two sets of oligos. One set contains a forward oligo that anneals downstream of modified positions, and the other set contains a different forward oligo that anneals upstream of the modified positions (Dong et al., 2012). Oligonucleotides used to assess modifications in U2 snRNA and 18S rRNA are depicted in Figures 3E and 4D, respectively. EpiScript RT was used on CMC treated and untreated U2 snRNA and AMV-RT on 18S rRNA.

### Site-directed cleavage of RNA by RNaseH

To test and quantify 2’-O-methylation levels of selected positions in U2, U5, U6 and U12 snRNAs we used a technique that utilizes RNAse H site-specific cleavage of RNA directed by RNA-DNA chimeric oligonucleotides. The method was developed in Joan Steitz’s laboratory and is based on the ability of 2’-O-methylated residues to protect RNA in RNA-DNA hybrid from RNase H digestion (Yu et al., 1997). It is known that in this assay the position of cleavage depends on the source of enzyme (Lapham et al., 1997). We tested RNase H (New England Biolabs) on control unmodified and single modified RNA oligos and designed experimental chimeric oligos accordingly. The chimeric oligos were the following:

U2-Gm25, rU_m_rG_m_rA_m_rU_m_**<u>dC</u>dTdTdA**rG_m_rC_m_rC_m_rA_m_rA_m_rA_m_rA_m_rG_m_

U2-Cm40, rG_m_rA_m_rA_m_rC_m_rA_m_**<u>dG</u>dAdTdA**rC_m_rU_m_rA_m_rC_m_rA_m_rC_m_rU_m_rU_m_ U5-Um41, rG_m_rU_m_rA_m_rA_m_**<u>dA</u>dAdGdG**rC_m_rG_m_rA_m_rA_m_rA_m_rG_m_rA_m_

U6-Cm77, rU_m_rG_m_rC_m_rG_m_rU_m_**<u>dG</u>dTdCdA**rU_m_rC_m_rC_m_rU_m_rU_m_rG_m_rC_m_

U12-Gm22, rU_m_rU_m_rU_m_rU_m_rC_m_**<u>dC</u>dTdTdA**rC_m_rU_m_rC_m_rA_m_rU_m_

In four-nucleotide DNA regions (boldface), the position that pairs with a 2’-O-methylated residue in tested RNA is underlined. RNA residues in the chimeras are 2’-O-methylated to stabilize oligos and increase specificity (rN_m_). Test RNAs in the amount of 3-5 µg were mixed with 20-50 pmol of a chimeric oligo in 15 µl of RNase H reaction buffer. The mixture was heated at 65o C for 5-10 min and annealed at 37o C for 10 min. Then 2-5 unites of RNase H (New England Biolabs) in 5 µl of 1x RNase H buffer were added to the annealed RNA-chimeric oligo mixture and the RNase H cleavage reaction was performed at 37o C for 1 h. In vitro transcribed (unmodified) snRNAs were used as controls for RNaseH digestion efficiency; RNA samples incubated without chimeric oligos or RNase H served as additional controls. The digested RNAs were separated on 8% polyacrylamide-8M urea gels, transferred onto a nylon membrane (Zeta Probe, Bio-Rad) and probed with digoxigenin (Dig)-labeled DNA fragments corresponding to human U2 [nt 61-3’ end], U5 [nt 1-41], U6 [nt 1-77], U12 [nt 23-3’ end]. Dig was detected using anti-Dig antibody conjugated with alkali phosphatase and CDP-Star substrate (Roche) according to manufacturer’s protocols. Li-Cor Odyssey Fc imaging system and Image Studio software were used to visualize and quantify results. Each RNA sample was assayed in 2-3 replicates; each replicate was split to run on two separate gels and to probe independently for reproducibility control.

## RT-PCR

Semiquantitative RT-PCR were performed on 1000ng of DNase RQ1 (Promega) treated total RNA using SuperScript III One-Step RT-PCR System (Thermo Fisher Scientific) following the manufacturer’s instructions. DNA fragments were separated on 4% for PCR products less than 100bp or 2% agarose gels and bands quantified using Image Studio Lite (LI-COR Biosciences). Primers (Thermo Fisher Scientific) used for RT-PCR are described in Supplemental Table S6.

### Electron microscopy

Monolayers of cells were fixed with 2.5% glutaraldehyde in 0.1 M sodium cacodylate buffer, postfixed with 1% osmium tetroxide followed by 2% uranyl acetate, and dehydrated through a graded series of ethanol, and the cells were lifted from the monolayer with propylene oxide and embedded as a loose pellet in LX112 resin (LADD Research Industries, Burlington, VT) in Eppendorf tubes. Ultrathin sections were cut on a Leica Ultracut UC7, stained with uranyl acetate followed by lead citrate, and viewed on a JEOL 1400 Plus transmission electron microscope at 80 kV.

## ACKNOWLEDGEMENTS

The microscopes used are maintained by the Einstein Analytical Imaging Facility (AIF), which is partially supported by the NIH funded Einstein Cancer Center (P30CA013330). For RNA FISH, access to the DS1 microscope was generously provided by Rob Singer. We are grateful to Louis Hodgson for the actin antibody. Leslie Gunther-Cummins and Xheni Nishku (AIF) prepared and imaged the samples on an electron microscope supported by an NIH funded Shared Instrumentation Grant (S10OD016214). This work was supported by grants from the National Institutes of Health, HL136662 (to U.T.M.) and R01 GM33397 (to J.G.G.). J.G.G. is American Cancer Society Professor of Developmental Genetics. Research in the lab of D.L.J.L. is supported by the Belgian Fonds de la Recherche Scientifique (F.R.S./FNRS), the Université Libre de Bruxelles (ULB), the European Joint Programme on Rare Diseases (‘RiboEurope’ and ‘DBAcure’), the Région Wallonne (SPW EER) (‘RIBOcancer’), the Internationale Brachet Stiftung, and the Epitran COST action (CA16120).

## SUPPLEMENTAL TABLES

**Table S1:**
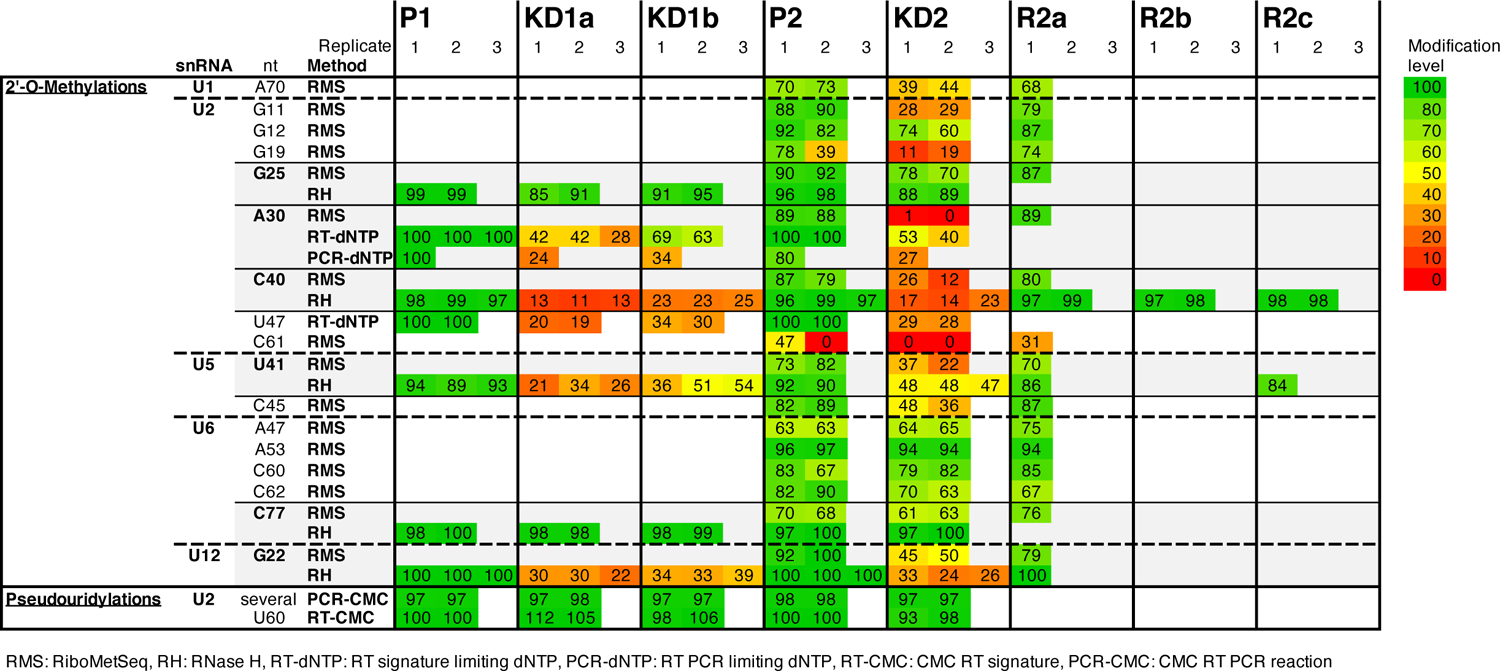
All snRNA modification scores assessed by RMS and individually

**Table S2.**
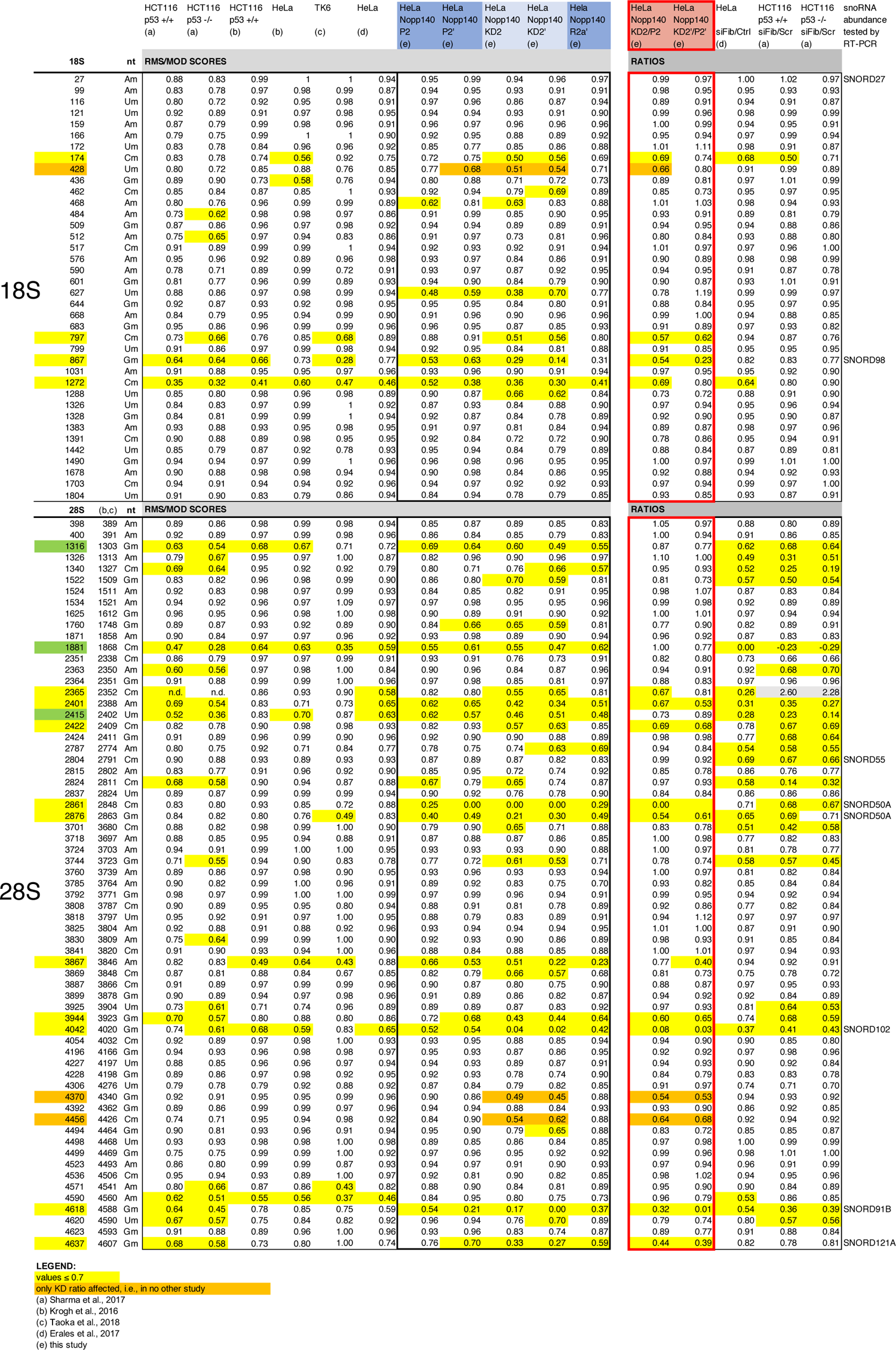
– RMS scores for ribosomal RNA compiled from available studies

**Table S3.**
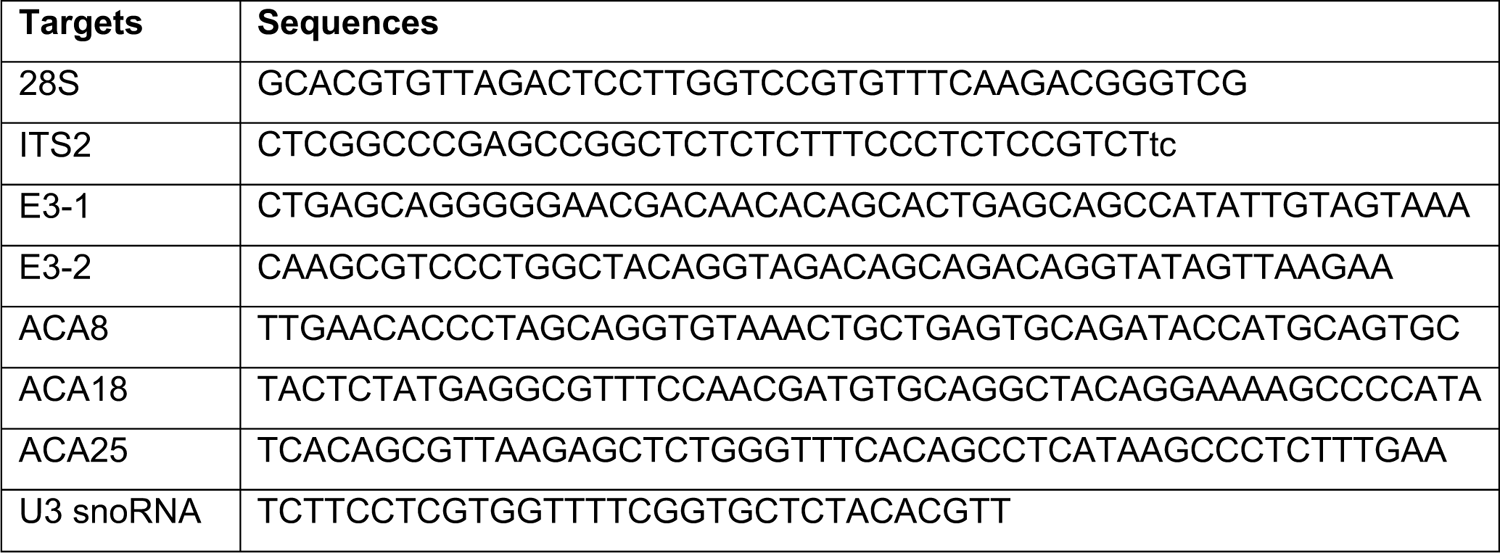
RNA FISH probes

**Table S4.**
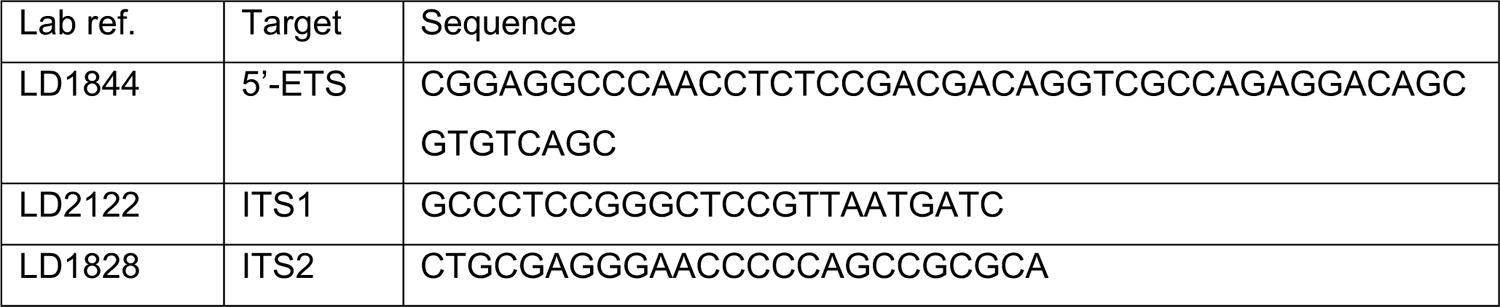
Primers used for pre-rRNA processing analysis

**Table S5.**
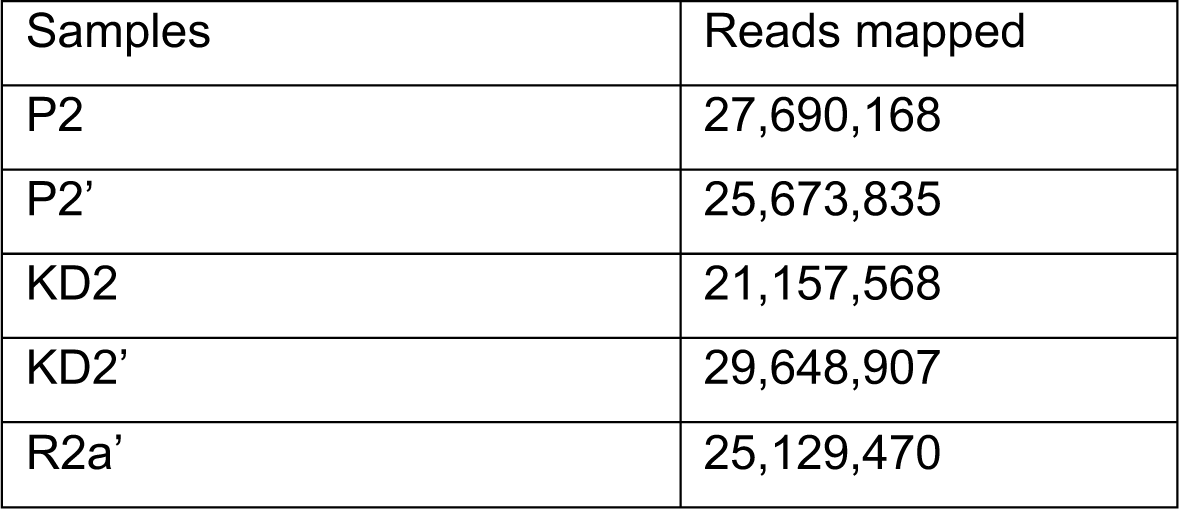
Details on sequencing for RiboMethSeq analysis

**Table S6.**
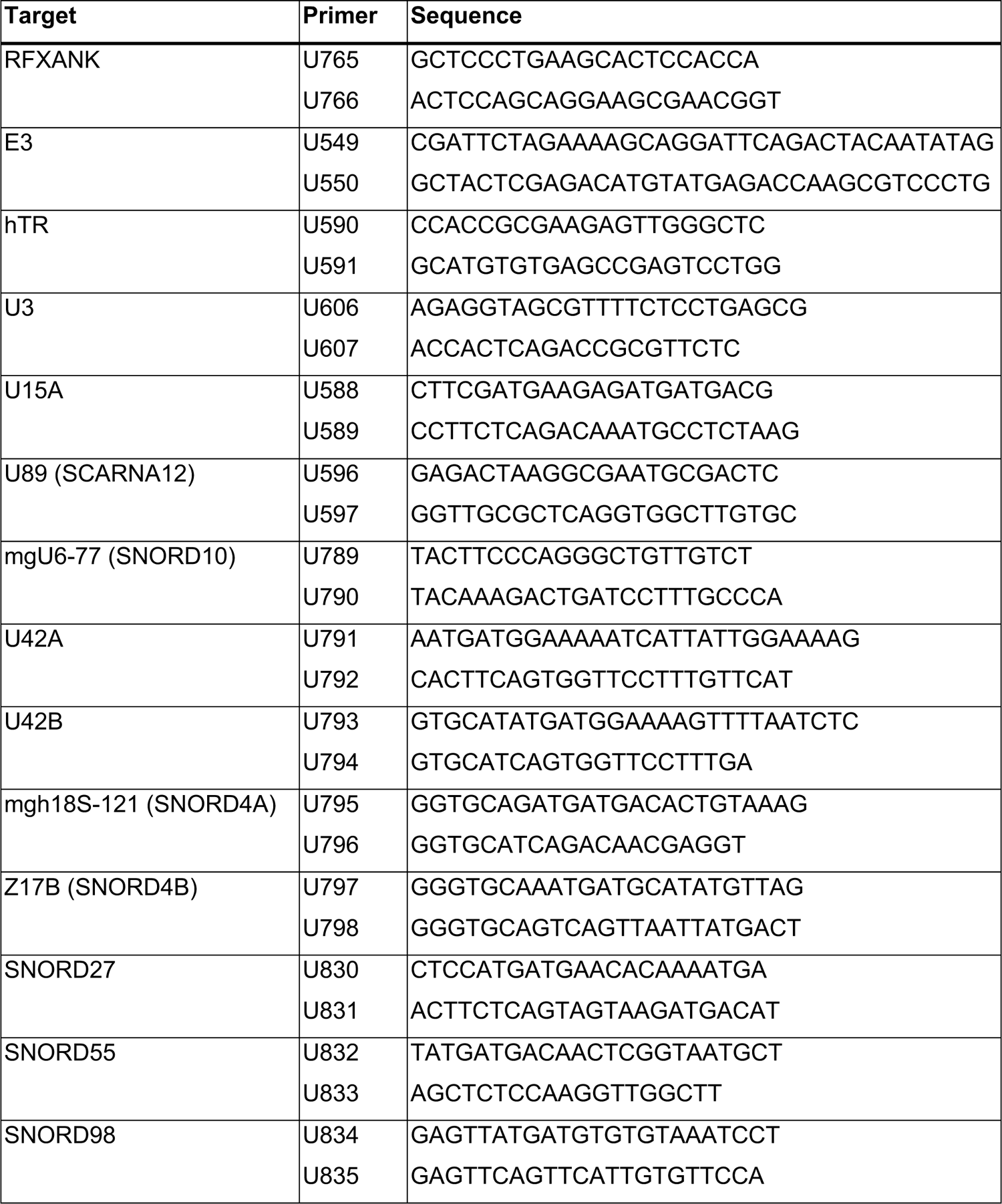
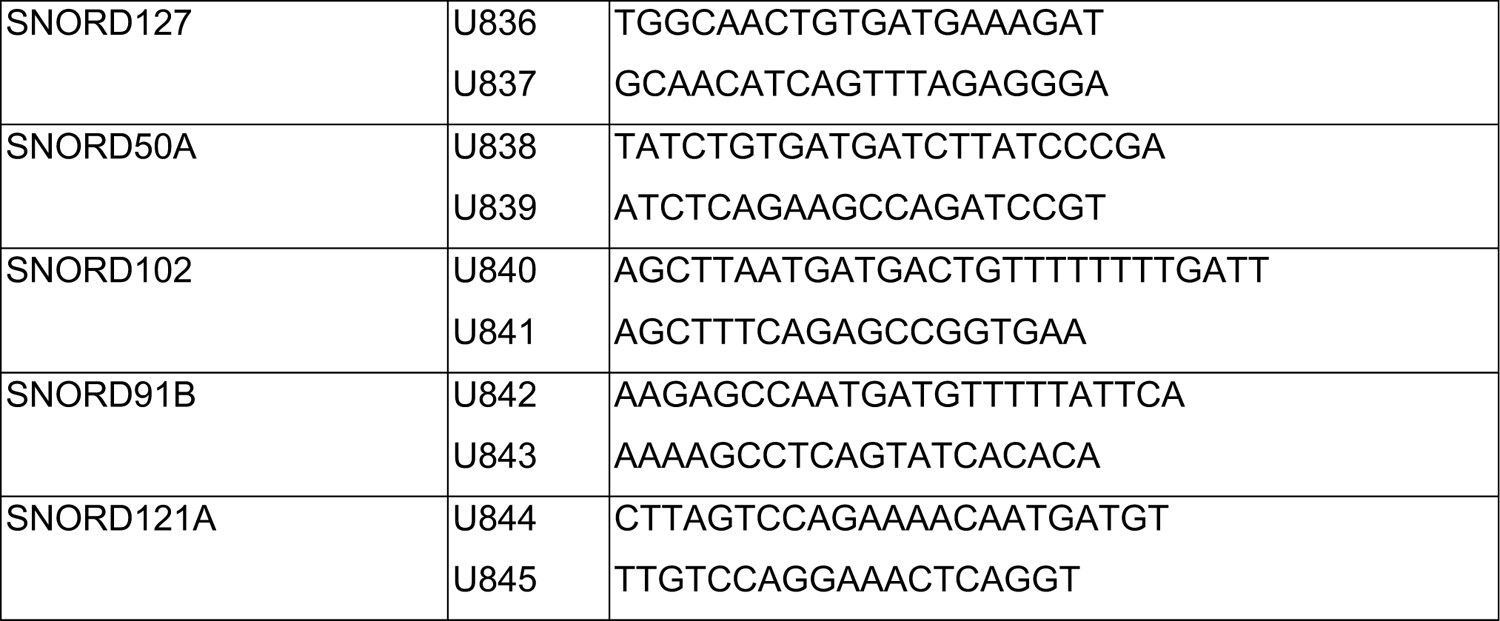
Primers for RT-PCR

## Notes

### Competing Interest Statement

The authors have declared no competing interest.

